# Folding of mRNA-DNA origami for controlled translation and viral vector packaging

**DOI:** 10.1101/2024.05.29.596417

**Authors:** Iris Seitz, Sharon Saarinen, Julia Wierzchowiecka, Jeroen J. L. M. Cornelissen, Veikko Linko, Mauri A. Kostiainen

## Abstract

mRNA is an important molecule in vaccine development and treatment of genetic disorders. Its capability to hybridize with DNA oligonucleotides in a programmable manner facilitates the formation of RNA-DNA origami structures, which can possess a well-defined morphology and serve as rigid supports for mRNA delivery. However, to date, comprehensive studies on the requirements for efficient folding of mRNA into distinct mRNA-DNA structures while preserving its translation functionality remain elusive. Here, we systematically investigate the impact of design parameters on the folding of protein-encoding mRNA into mRNA-DNA origami structures and demonstrate the importance of the availability of ribosome-binding sequences on the translation efficiency. Furthermore, these hybrid structures can be encapsulated inside virus capsids for protecting them against nuclease degradation and also for enhancing their cellular uptake. This multicomponent system therefore showcases a modular and versatile nanocarrier. Our work provides valuable insight into the design of mRNA-DNA origami structures contributing to the development of mRNA-based gene delivery platforms.

## 1 Introduction

Nucleic acids play a key role as therapeutic agents by directly targeting genes and subsequently modulating their expression [1]. Therefore, they find versatile applications in protein replacement therapy [2], gene silencing [3] and as vaccines [4, 5]. Nucleic acid therapeutics are highly specific [1], however, their success is dependent on the (intra)cellular delivery, which can be enhanced by complexing the nucleic acids into nanoscale particles. The transfection efficiency is furthermore influenced by the size, shape, and the surface chemistry of the nanoparticle [6, 7]. Subsequently, packaging of high molecular weight nucleic acids, including gene-encoding nucleic acids and mRNAs, into a compact, predefined shape would be desirable. However, this remains challenging using conventional agents such as lipids [8].

An attractive alternative is offered by utilizing the DNA origami technique [9]. It allows the fabrication of predefined, custom-shaped two- and three-dimensional DNA nanostructures [10, 11] by folding a long, single-stranded DNA (ssDNA) scaffold into a desired shape with the help of ”stapling” DNA oligonucleotides [9, 12]. DNA origami structures are highly addressable, allowing for site-specific functionalization [13] with targeting agents like antibodies, affibodies, aptamers and peptides for improved drug delivery [14]. The choice of scaffold strand is no longer limited to generic, bacteriophage-genome-derived sequences [15], as the fabrication schemes based on synthetic, gene-encoding scaffolds have become available [16, 17]. Similarly, the DNA scaffold can be replaced with mRNA, which facilitates protein translation without the need to enter the nucleus [18]. The mRNA can be folded into the predefined shape [19] using either kissing loops following the RNA origami technique [20–22] or ssDNA staple strands, resulting in a mRNA-DNA structure. Recently, several RNA-DNA origami structures have been presented, aiming to mediate the delivery of both mRNA and antisense oligonucleotides [23–26].

The design of (multilayer) DNA origami structures is commonly based on a square or a honeycomb lattice, resembling the canonical *B* -form of duplex DNA with 10.67 or 10.5 base pairs (bp) per helical turn, respectively [27]. In contrast, in RNA origami, the double-helical RNA domains adopt the *A*-form, resulting in 11 bp per turn [21]. Previously reported RNA-DNA origami were designed to have either 10.5, 10.67, or 11 bp per turn [23, 25, 28]. Although a tendency towards the *A*-form, *i.e.*, 11 bp per turn has been suggested for short RNA-DNA structures [29], a thorough study on the design parameters for folding and subsequent translation has not yet been realized.

In this article, we explore the design of a six-helix bundle (6HB) mRNA-DNA origami with respect to its folding and translation properties. To this end, a protein-encoding mRNA scaffold is folded into distinct nanostructures using short ssDNA staples (Figure 1a). Thus, certain parts of the mRNA, for instance the poly(A)-tail, can be purposely left non-hybridized. First, we investigate the impact of the internal design, *i.e.*, the crossover (CO) density and the number of bp per helical turn, on the folding efficiency, while also changing the buffer environment and temperature gradient of the folding reaction (Figure 1a,b). By additionally adjusting the single-stranded parts in the mRNA-DNA origami, we ensure the preservation of the innate translation property of mRNA once it is folded. To reduce the susceptibility of nucleic acid structures towards nucleases [30], we use virus capsid proteins to encapsulate the mRNA-DNA origami, resulting in a structurally well-defined protein-based nanocarrier. The protein capsid is not only found to improve protection, but it also enhances the nanocarrier’s uptake into cells and henceforth the mRNA’s translation into the target protein while exhibiting minimal toxicity in the system studied.

**Fig. 1.**
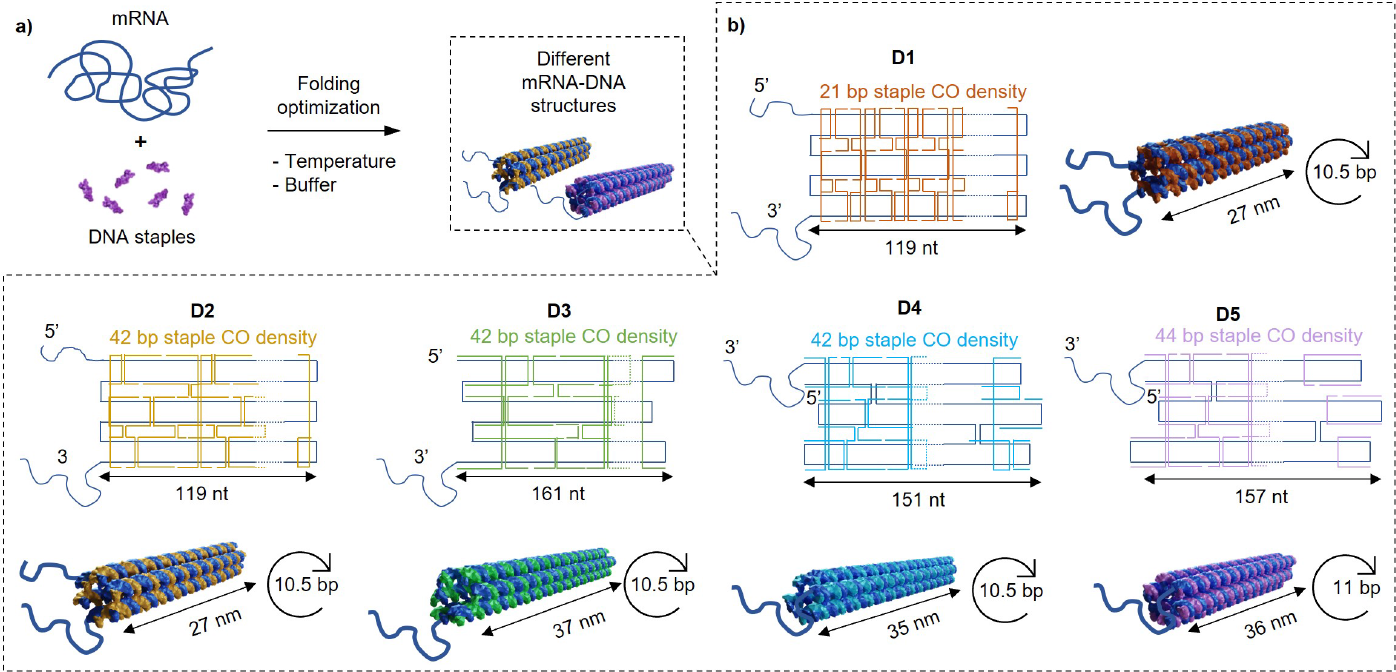
**a**, A 996-nt mRNA including an EGFP-encoding sequence is hybridized with short ssDNA strands by thermal annealing resulting in mRNA-DNA origami variants which dif- fer in their internal design. The folding conditions with regard to buffer composition and annealing temperature are optimized for every design. **b**, The designs (D1–D5) differ in the scaffold routing with respect to the scaffold crossovers (CO) and embedment of the 5’-cap region, the staple CO density (orange, yellow, green, light blue, light purple), and the heli- cal rise of the mRNA-DNA duplex, given as bp per turn.

## 2 Results

### 2.1 Design rationale for mRNA-DNA origami

We designed five different honeycomb lattice -based 6HB mRNA-DNA origami variants (Figure 1b) to investigate possible structural limitations during the folding process as well as their properties regarding stability and translation efficiency. To facilitate the translation characterization, we selected a 996- nucleotide (nt) long mRNA, which includes an enhanced green fluorescent protein (EGFP) encoding sequence, as our scaffold (blue).

For folding the mRNA scaffold with DNA staple strands, we started off with two variants, where only the mRNA’s EGFP encoding part (714 nt) was hybridized, *i.e.*, the poly(A)-tail and the 5’-cap region of the mRNA remained single-stranded, thus resulting in 27-nm long structures. These variants additionally featured 7–8-nt long scaffold loops, *i.e.*, single-stranded regions of the scaffold at the both ends. Since the density of inter-helical staple COs between neighboring helices was previously found to affect the stability of 6HB DNA origami structures [31], two different CO densities were tested: the COs were spaced either at 21 bp (Figure 1b, D1, orange, top right) or 42 bp intervals (D2, yellow, bottom left) (*i.e.*, 21 bp or 42 bp between COs that link the same neighboring helices).

Keeping the CO density at 42 bp, a 37-nm long version without the scaffold loops was designed, in which the 5’-cap region was hybridized with the staples (D3, green, bottom middle-left). To further increase the stability and rigidity, scaffold COs were included in the structures to obtain a structure with 35 nm in length (D4, light blue, bottom middle-right).

The helical rise of the variants D1–4 was set to 10.5 bp per turn, thus resembling *B* -DNA and being in line with previously reported RNA-DNA origami [23, 25]. To decrease strain in the origami structure, complying with the tendency of short RNA-DNA duplexes towards the *A*-DNA geometry [29], the design of the fifth variant (36 nm in length) also featured two scaffold crossovers and the hybridization of the 5’-cap region, but its helical pitch was set to 11 bp per turn (D5, light purple, bottom right).

In addition, all design variants included four staples that could be replaced with extended staples to further facilitate attachment of ATTO590 dye (A590) -modified DNA stands through DNA-DNA hybridization. It allows tracking of the origami, in addition to ethidiumbromide (EtBr), under red light (Alexa A647 channel for gels and AF594 channel for uptake into cells).

### 2.2 Optimization of the folding environment for mRNA-DNA origami

The folding of DNA origami structures is usually performed in a buffered one-pot reaction by first heating the solution and then gradually cooling it to room temperature. The optimal buffer composition (folding buffer; FOB), including cations to screen the negative charges of the nucleic acid backbones as well as the temperature gradient are both structure specific [27]. Since commonly employed Mg^2+^ ions can degrade RNA at high temperatures [25], a Tris-ethylenediaminetetraacetic acid (EDTA) -based buffer (TE) supple- mented with 0–100 mM NaCl was initially chosen for folding the mRNA-DNA origami. The reaction mixture was cooled down overnight from 65 °C to 20 °C (Figure 2a-d, Note S1).

**Fig. 2.**
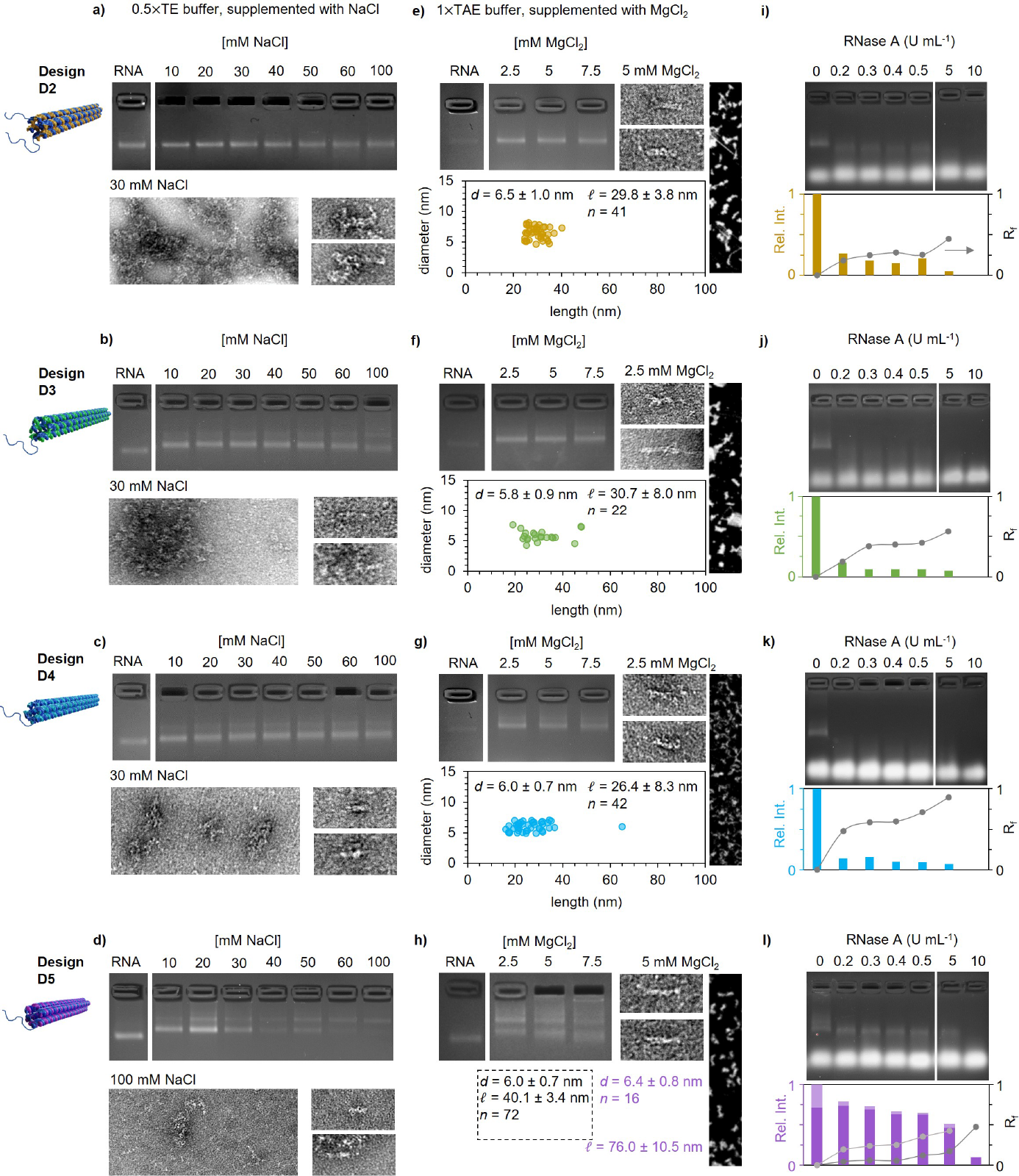
**a–d**, Optimization of the buffer conditions in the folding reaction for each design variant by supplementing 0.5*×*TE buffer with 0–100 mM NaCl. The success of the folding reaction is evaluated from the change in electrophoretic mobility on a 3.5 % agarose gel (top). Negative-stain TEM images (bottom) display aggregated structures (left; image width corresponds to 325 nm) as well as discrete folded structures (right, image width corresponds to 100 nm). **e–h**, Optimization of the folding conditions using isothermal temperature and screening of the MgCl2 concentration in 1*×*TAE buffer. Analysis of the optimized conditions by assessing the electrophoretic mobility of the folded structures (top left), negative-stain TEM images (top middle; image width corresponds to 100 nm) and AFM images (top right; image width corresponds to 200 nm) of plain mRNA-DNA origami. Observed dimensions of the folded structures (bottom). For D5, the average dimensions are calculated separately for monomers (dark) and dimers (light). **i–l**, Evaluation of the folding yield RNA-DNA origami obtained under optimized conditions by exposure to RNase A in a high salt buffer. Both the intensity (bars) and the electrophoretic mobility (*R_f_* , grey) from a 2 % agarose gel (top) are analyzed with respect to the untreated sample (bottom).

The outcome of the folding reactions was monitored using agarose gel electrophoresis (AGE), transmission electron microscopy (TEM), and atomic force microscopy (AFM). Both D1 (Figure S1a) and D2 (Figure 2a) showed a very similar electrophoretic mobility compared to the mRNA scaffold for all NaCl concentrations tested (top panel). At high salt concentrations a decrease in the intensity and broadening of the leading band was observed. However, the folding efficiency was low, since mainly unfolded or aggregated mRNA were observed under TEM (30 mM NaCl; Figure 2a, bottom panel, Figure S2a) and the few folded structures were shorter than expected (ca. 21 nm).

Similar results were also obtained for D3 and D4. While D3 and D4 (Figure 2b,c, top panel) showed reduced mobility compared to the plain mRNA scaffold, also an additional, even slower migrating band appeared. The number of correctly folded structures observed in TEM was low for both D3 and D4 (30 mM NaCl; Figure 2b,c bottom panel, Figure S2b,c) as the lengths of the folded objects appeared to be only ca. 16–29 nm (D3) and 20–29 nm (D4). The low yield and observation of only truncated structures for D2–D4 suggests a potential kinetic trap during the origami folding process.

By increasing the helical pitch from 10.5 bp per turn to 11 bp per turn in D5, two bands with decreased mobilities were observed (Figure 2d, top). In comparison to D4, it was noticeable that the leading band vanished gradually at higher NaCl concentration. When folded with a buffer supplemented with 30 mM NaCl, a higher number of folded structures compared to D4 was observed, however, the large spread in the size distribution remained, with the majority of the structures having lengths between 16–26 nm, suggesting the presence of partially folded structures (Figure S3a). An increase of NaCl to 100 mM resulted in an overall increase in the length of the observed folding products, displaying also a low number of structures with the designed dimensions of D5 (Figure 2d, bottom, Figure S3b).

Aiming to further increase the yield of the folding reactions for all structure variants, the temperature gradient was changed to a short (15 min) isothermal folding at 55 °C, and therefore, also the buffer composition could be altered to a Tris-acetate-EDTA (TAE) buffer supplemented with MgCl_2_ (Figure 2e- h). These changes resulted in better folding outcomes for all our designs. The shift in electrophoretic mobility compared to plain mRNA was still minimal for D1 (Figure S1b,c) and D2 (Figure 2e, top panel), but TEM and AFM showed correctly folded D2 variants (Figure 2e, right panel, Figure S4a). The observed D2 dimensions (given as average (avg) *±* s.d. throughout) both in length (*l* ) with *l*_avg_ = 29.8 *±* 3.8 nm and diameter (*d* ) with *d*_avg_ = 6.5 *±* 1.0 nm (Figure 2e,bottom) correspond well with the designed ones.

For both D3 and D4 (Figure 2f,g, and Figure S4b,c, respectively) full length structures could be observed, however, the scattering in the length distribution persisted also in these conditions, resulting in *l*_avg_ = 30.7 *±* 8.0 nm for D3 and *l*_avg_ = 26.3 *±* 8.3 nm for D4.

Analysis of the folding reaction of D5 on an agarose gel (Figure 2h, Figure S4d) revealed a leading band, which migrated slower than the plain mRNA, and an additional band, suggesting the formation of dimers. Indeed, both monomers (*l*_avg_ = 40.1 *±* 3.4 nm, *d*_avg_ = 6.0 *±* 0.9 nm) and dimers (*l*_avg_ = 76.0 *±* 10.5 nm, *d*_avg_ = 6.4 *±* 0.8 nm) were observed under TEM, and their dimensions were in good agreement with the designed ones.

Despite the observation of full length structures for all mRNA-DNA origami variants, the folding yields were found to differ between the designs. To further quantify the success of the folding reaction, the structures were folded in their optimized conditions, *i.e.*, 1*×*TAE supplemented with 5 mM MgCl_2_ for D1, D2 and D5, and 2.5 mM MgCl_2_ for D3 and D4, and subsequently, they were subjected to treatment with RNase A (0–10 U mL*^−^*^1^, Figure 2i-l), since it can digest single-stranded RNA (ssRNA) [32] at high salt concentrations. In this context, the correctly folded variants D3–D5 contained ssRNA only in the form of the poly(A)-tail, while all unfolded, truncated variants or aggregates could also display (partially) non-hybridized mRNA to various extent. In addition, variants D1 and D2 contained ssRNA scaffold loops, and their 5’-cap region remained non-hybridized, however, the capping of the mRNA (see Methods, Section 3.1) should enhance its overall stability [33].

Possible degradation of the origami variants upon RNase A addition was monitored by comparing the relative intensities of the leading bands and the relative retention factors (*R_f_* , see Methods), *i.e.*, the difference between the relative migration distance of the treated sample and that of the untreated origami (0 U mL*^−^*^1^, first lane in each gel). While the plain mRNA scaffold was digested immediately (although used at three times as high concentration as the origami (Figure S5)), all folded variants except D1 (Figure S1d) elicited an increased resistance.

For all variants D2–D5 (Figure 2i-l), both a decrease in the intensity of the leading band and an increase in *R_f_* were observed when incubated with 0.2–0.5 U mL*^−^*^1^ of RNase A. The most significant changes upon treatment with 0.2 U mL*^−^*^1^ were detected for D4, with a leading band intensity decrease of 86 % as well as an increase in *R_f_* to 0.48, thus indicating the largely exposed mRNA and a relatively low folding yield. The digestion of D3 was slower than that of D4, however, upon treatment with 0.3 U mL*^−^*^1^ the *R_f_* had increased to 0.38. Interestingly, both the *R_f_*and the leading band intensity showed a plateauing behavior for D2–D5, which reflects the folded, double-stranded segments of the origami, before fully vanishing when increasing the amount of RNase A drastically to *≥* 5 U mL*^−^*^1^.

Overall, the differences in the observed folding yields for origami variants can be attributed to the internal design. D2 has no scaffold COs, and the staple CO density of 42 bp makes the structure rather flexible allowing for mechanically less restrained folding and also high yields. In comparison, an increase of the CO density in D1 (21 bp intervals) seems to hamper its folding, since almost no folded structures could be observed.

Despite having a staple CO density of 42 bp in D3, it differs from D2 in the scaffold routing, including the removal of scaffold loops and hybridization of the 5’-cap region, alterations which seem to decrease the folding yield. The addition of two scaffold crossovers in the scaffold routing of D4, in combination with the mismatch in the helical pitch between the mRNA scaffold and the DNA staples most likely reduced the degree of freedom for the staple strands to bind correctly, thus resulting in a further decrease of the folding yield.

Increasing the helical pitch from 10.5 bp to 11 bp per turn in D5 resulted in a high yield of well-folded structures. However, D5 tended to form dimers, and therefore, the folding conditions were further optimized (Figure S6), leading to a final buffer composition of 1*×*TAE supplemented with 5 mM MgCl_2_ and 1 mM NaCl. In addition, folding conditions reported for a wireframe RNA-DNA origami structures (10 mM HEPES (4-(2-hydroxyethyl)piperazine- 1-ethane-sulfonic acid) buffers supplemented with KCl) [26] were also tested, however, these conditions resulted in very similar folding yields as observed in optimized TAE -based buffers for all mRNA-DNA origami variants (Note S2).

### 2.3 Importance of the ribosome binding site positioning in translation

Our results above show that the folding yield of each mRNA-DNA origami variant is dependent on the choice of the buffer and the thermal gradient. From the tested variants, D2 and D5 were found to be the most versatile ones as they folded successfully in various conditions regarding both the buffer composition and the temperature. Therefore, these variants were selected for the further studies.

For potential applications, also the disassembly of the mRNA-DNA origami and subsequent translation of the mRNA are highly important. Therefore, the translation of purified D2 and D5 was first studied in an extra- cellular environment using reticulocyte lysate (Figure 3a), and the outcome was analyzed by native polyacrylamide gel electrophoresis (PAGE). While D2 was successfully translated into EGFP (Figure 3b,c, first lane, yellow bar), appearing as a fluorescent band upon excitation with blue light (Alexa A488 channel), no signal was obtained for D5 (second lane, light purple bar).

**Fig. 3.**
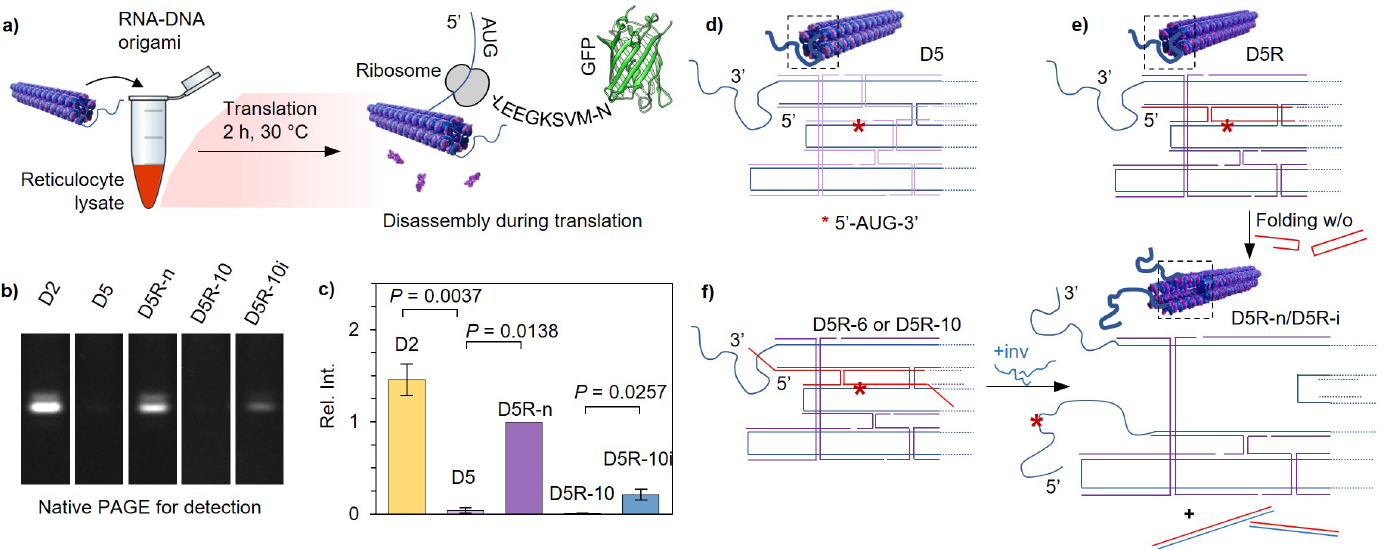
**a**, Addition of the folded mRNA-DNA origami to reticulocyte lysate results in extracellular translation into EGFP. **b**, The fluorescence signal is detected using native PAGE. **c**, Relative intensities of the fluorescence signals obtained from (b). **d**, Design of the staple strands (light purple) that hybridize with the 5’-cap region of the mRNA for D5, with the position of the start codon in mRNA scaffold (blue) being marked by an asterisk. **e**, Redesign of the staples leaves only two replacement strands that hybridize with the 5’-cap region of the mRNA (variant D5R, replacement strands marked with red). **f**, The replacement strands can also be designed with 3’-overhangs (D5R-6 or D5R-10). The addition of invader strands (inv, blue) leads to hybridization with the replacement strand, which subsequently releases the 5’-cap region of the mRNA. The same result can be obtained from (e) by omitting the replacement strands in the folding mixture.

In eukaryotes, the process of ribosome recruitment starts usually at the 5’- cap region of the mRNA. As soon as the start codon is reached, translation is initiated [34]. Apart from the helical geometry, D2 and D5 differ mainly in the configuration of the 5’-cap region. While it is part of the folded core of D5 (Figure 3d, the start codon marked with an asterisk), it is non-hybridized and therefore freely available in D2.

To probe the role of hybridization of the mRNA’s 5’-region, the staple design downstream of the 5’-cap of D5 was revised. In the revised version (D5R, purple), the 5’-region of the mRNA was hybridized only with two replacement staple strands (red strands in Figure 3e). Both replacement staple strands were also designed with 3’-overhangs (variant D5R-6 or D5R-10, Figur 3f, Figure S11), thus facilitating a triggered release of the replacement strands through a toehold-mediated strand displacement reaction using invader strands (inv, blue). Now, by either omitting these two replacement staple strands from the staple mixture or by adding the invader strands, a freed 5’-cap region was obtained (D5R-n or D5R-i).

The importance of the accessibility of the 5’-cap region was confirmed by reperforming the translation studies with D5R-n, which revealed successful mRNA translation into EGFP (Figure 3b,c third lane, purple bar). However, in comparison to D2, the protein translation yield was reduced by one third. Additionally, the irreversible unfolding of the 5’-cap region through the strand displacement reaction and the subsequent translation of the mRNA was demonstrated. To this end, the translation of D5R-6 and D5R-10, which featured 6 nt or 10 nt long toeholds on the replacement strands, respectively, was investigated first without the addition of an invader strand. As expected, no translation activity was observed (Figure 3b,c, fourth lane, orange bar, Figure S12a), confirming that the toeholds do not interfere with the origami folding. Subsequently, D5R-6 and D5R-10 were incubated with 10*×* excess of the corresponding invader strands for 10 min prior to their addition to the reticulocyte lysate. While the signal obtained from D5R-6i was negligible (Figure S12a), a weak fluorescence band was observed for D5R-10i (Figure 3b,c, fifth lane, blue bar), showing that the toehold-mediated irreversible unfolding of the 5’-cap region allowed to recover 20 % of the fluorescence intensity when compared to D5R-n.

To rule out false positive signals originating from the unfolded mRNA that could have been left in the solution after the folding reaction and purification, the translation capabilities of all mRNA-DNA origami variants without a purification step were similarly tested (Figure S12b). The fluorescence intensities from D2, D5 and D5R-n corresponded to the previously observed results, with D2 showing the highest translation activity while the translation of D5 was prohibited. A fluorescence signal could also be detected for D3, which might have been expected based on the folding evaluation. The majority of the observed D3 variants were truncated under TEM, which indicates that the structures were only partially folded, and thus contained longer segments of free, unhybridized mRNA than intended by the design. Therefore, it is also possible that the start codon in D3 was exposed. In comparison, the translation of D4 was at the detection limit, suggesting a correctly folded 5’-cap region. Surprisingly, the EGFP signal obtained for D1 exceeded those of all other variants, suggesting mainly aggregation during the folding reaction, and therefore resulting in easy accessibility of the start codon.

### 2.4 Protection of mRNA-DNA origami using viral capsids

Nucleic acids are known for their susceptibility to degradation by nucleases, making their therapeutic use challenging. To maintain the intactness of nucleic acid nanostructures, several strategies have been developed [14, 30]. One feasible approach is to create a protective shell around the nucleic acid structures [35, 36]. To achieve this, it has been previously demonstrated that capsid proteins isolated from cowpea chlorotic mottle virus (CCMV) can reassemble on DNA origami structures by exploiting electrostatic interactions between the negatively charged origami and the positively charged *N* -terminus of the capsid protein [37, 38].

To assess the complexation between the capsid proteins and the mRNA-DNA origami, D2 and D5R-n were mixed with increasing concentrations of CCMV capsid proteins and the shift in the electrophoretic mobility of the formed complexes was monitored using AGE (Figure 4a for D5R-n and Figure S13a for D2). A significant decrease in the mobility was observed for both origami variants already at a protein excess (given as c_capsids_/c_origami_) of 100, and increasing the protein concentration from that led to further decrease in the mobility.

**Fig. 4.**
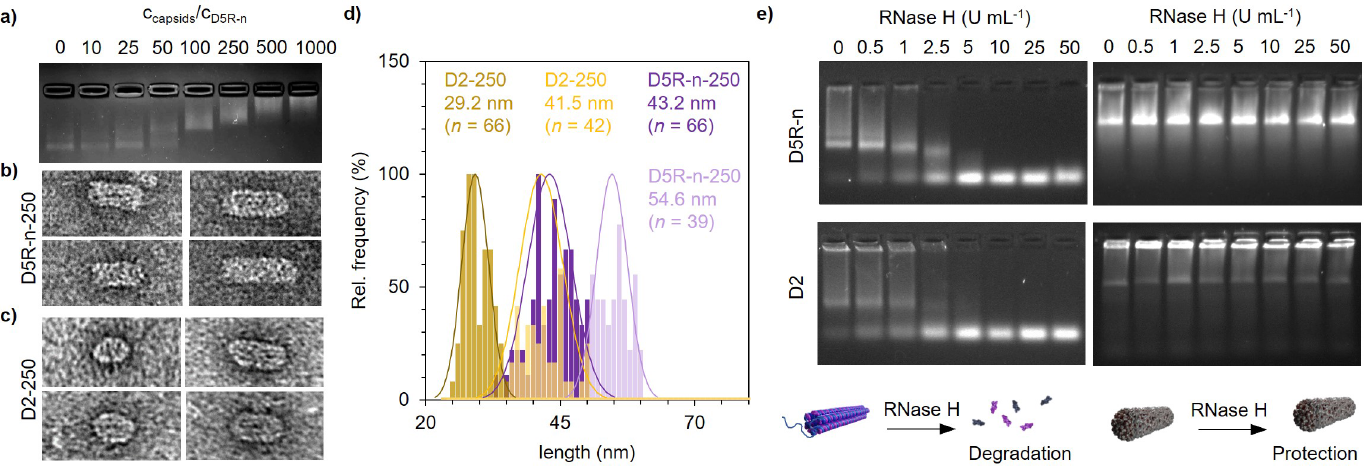
**a**, The electrophoretic mobility of D5R-n changes with increasing virus capsid protein concentration. **b, c**, Negative-stain TEM images showing the development of a single protein layer on D5R-n (b) and D2 (c) when using a protein excess of 250. The obtained complexes are grouped based on their lengths; the image width corresponds to 100 nm. **d**, Observed length distributions for D2 and D5R-n -templated complex populations with 250*×* excess of the virus capsid proteins and the calculated Gaussian distribution (solid line). **e**, RNase H digestion of plain (left) and complexed (right) D5R-n (top) and D2 (bottom).

Aiming to use as low protein concentration as possible for the coating, mRNA-DNA origami complexed with an excess of 250 were selected and imaged with TEM. For both origami variants, the formation of a highly ordered protein coat was observed (Figure 4b,c, Figure S13b,c). This can be also clearly seen from the increased diameter of the imaged objects: for D5R-n from *d*_avg_ = 6.0 *±* 0.9 nm (plain origami) to *d*_avg_ = 18.7 *±* 1.5 nm (complex) and for D2 from *d*_avg_ = 6.5 *±* 1.0 nm (plain origami) to *d*_avg_ = 19.2 *±* 2.0 nm (complex). The dimensions are in line with previously reported DNA origami-capsid protein complexes [37].

Interestingly, the coated structures showed a heterogeneity in length, which could be grouped into two main populations (Figure 4d). The first population is formed when the proteins coat the folded core of the structure resulting in complexes with lengths of *l*_avg_ = 29.2 *±* 2.5 nm (D2, dark yellow) and *l*_avg_ = 43.2 *±* 4.1 nm (D5R-n, dark purple). The second population consists of complexes with *l*_avg_ = 41.5 *±* 4.0 nm (D2, light yellow) and *l*_avg_ = 54.6 *±* 2.9 nm (D5R-n, light purple), respectively, suggesting that the free 5’-cap region together with the poly(A)-tail can further template the capsid protein shell formation (Figure S13d).

To test the possible protection of the coating, both the uncoated and complexed structures were incubated with RNase H (0–50 U mL*^−^*^1^) (Figure 4e), which is known to attack RNA in RNA-DNA complexes [39]. The protein shell was found to enhance the stability of both D5R-n (top) and D2 (bottom) when treated with RNase H, making it possible for the complexed structures to withstand nuclease concentrations up to 50 U mL*^−^*^1^ (right gels). The uncoated, plain variants started to degrade already at 0.5 U mL*^−^*^1^, and only subtle differences in the stability and digestion profiles between the variants were observed (Figure 4e, left gels).

In addition, the susceptibility of the structures towards DNase I was studied (Figure S14). DNase I is known to act non-specifically on ssDNA and the minor groove of double-stranded DNA (dsDNA) by cleaving the phosphodiester bonds [40]. Despite its preference for dsDNA, the enzyme may also show some residual activity toward RNA-DNA duplexes [40]. Throughout the tested DNase I concentration range (0–50 U mL*^−^*^1^), the electrophoretic mobility of D5R-n remained constant, suggesting that D5R-n can withstand the enzyme treatment. For D2, in contrast, an increase in smear was observed once it was treated with 0.5 U mL*^−^*^1^ DNase I, most likely originating from the degradation of the DNA staples.

Since both variants were equipped with A590-modified DNA strands through hybridization to complementary DNA staple overhangs (*i.e.*, DNA-DNA duplex formation), the digestion processes could additionally be monitored in the fluorophore channel (Figure S14, Alexa647, A647). For both variants, a release of the fluorophores was detected once DNase I was added. However, the leading band of D5R-n could be observed over a wider concentration range in the A647 channel, suggesting a slower degradation of the fluorophore-containing strands. Nevertheless, the virus capsid coating efficiently protected the whole origami from degradation with up to 50 U mL*^−^*^1^.

### 2.5 Transfection and translation *in vitro*

Having ensured the accessibility of the 5’-cap region of the mRNA for translation and established a protein shell to prevent degradation by nucleases, the uptake efficiency of the mRNA-DNA origami was tested *in vitro* using HeLa cells (Figure 5, Figures S15-18). Briefly, the cells were seeded into a 96-well plate and grown for 24 h. Subsequently, the media was exchanged to Opti-MEM, and the cells were incubated with the samples for 16 h or 24 h. We used A590 to track the uptake of D2 and D5R-n into the cells, and the signal read-out was performed in 1*×*PBS using a microplate reader (excitation/emission of 490/520 nm for EGFP and 584/616 nm for A590) and fluorescence microscopy (GFP filter at 470 nm for EGFP and AF594 filter at 590 nm for A590).

**Fig. 5.**
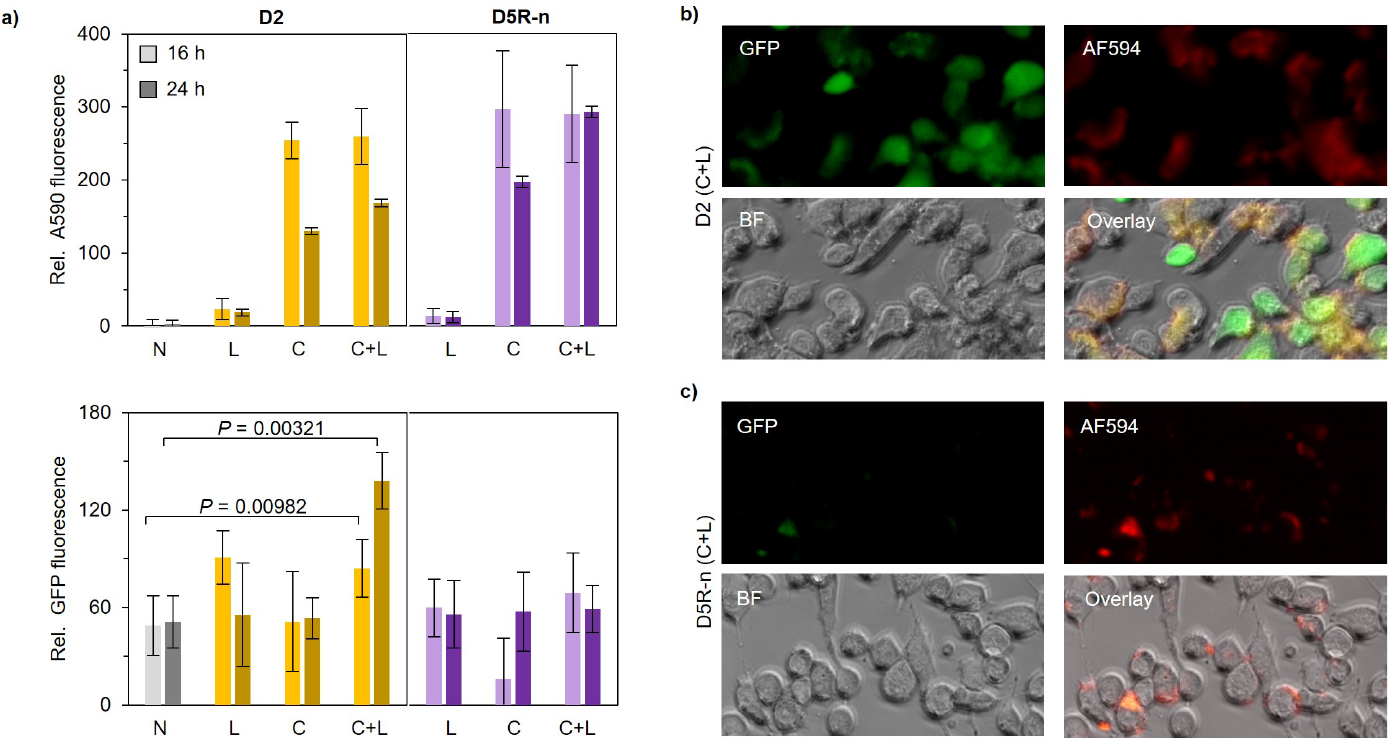
**a**, Monitoring the uptake of naked (N) and D2 (yellow) and D5R-n (purple) complexed with either LP2000 (L), CCMV capsid proteins (C) or a combination of capsid proteins and LP2000 (C+L) into HeLa cells (top) and the translation activity of the same samples (bottom) for 16 h (light) and 24 h (dark). The data is given as avg *±* s.d. of *n* = 3 samples. **b,c**, Microscopy images recorded from the GFP, AF594 and bright field (BF) channels, showing the translation of D2 and D5R-n, respectively. Both samples were transfected with C+L for 24 h; the image width corresponds to 200 µm.

We started out with investigating the transfection efficiency of naked structures (N) and structures complexed with virus capsid proteins (C, protein excess of 250) (Figure 5a, top). In addition, these samples were compared to a control sample with the commercially available transfection agent Lipofectamine 2000 (LP2000, Note S8). While the uptake of both naked (N) and LP2000 treated (L) structures (Figure 5a, top) was found to be low, the A590 signal increased drastically once the viral capsid coating was applied onto the structures (C). This trend was observed for both 16 h and 24 h sample incubation with the cells, however, a higher fluorescence signal was obtained after 16 h. Moreover, a sequential combination of virus capsids and LP2000 was tested (C+L), in which mRNA-DNA origami was first complexed with the capsid proteins, followed by the addition of the LP2000 (Figure S19d). Interestingly, the uptake of such complexes was comparable to the C-samples, suggesting the virus capsid proteins being the driving force in the enhanced uptake. Notably, higher transfection efficiency were found for D5R-n for both C and C+L samples in comparison to D2.

Despite the high internalization of the capsid-coated mRNA-DNA origami (C and C+L), the EGFP signal (Figure 5a, bottom) from the C-samples was similar as for the naked structures, suggesting the inaccessibility of the mRNA for the ribosomes. In comparison, L-samples were found to elicit a similar level of translation. Only in the presence of both virus capsid proteins and LP2000 (C+L), a significant change in the EGFP signal was observed, being more pronounced after 24 h. Moreover, it was noted that the average EGFP intensity for D2 was higher than for D5R-n, but it was also expected based on the observation during the extracellular transcription. As a control, plain mRNA was subjected to transfection using the same conditions (Note S9). Here, the EGFP signal was found to decrease over time, being highest after 4 h (Figure S21a), indicating that the process of decomplexing the mRNA-DNA origami-based nanocarrier is relatively slow.

Finally, the toxicity of the samples towards the cells was evaluated (Supplementary Note S10). While the cell viability decreased by approximately 50 % upon treatment with L-samples, naked structures and samples coated with the virus capsids (C) or with the combination of C+L were not cytotoxic within the studied application relevant concentration range.

### 2.6 Discussion

We have investigated the folding behavior of a short, 6HB mRNA-DNA origami structure with respect to preserving the mRNA’s innate functionality, *i.e.*, its ability to be translated into a protein. To this end, we initially drafted five origami variants, which differed in their internal design. Commonly, the folding of DNA origami is considered to depend on the structure’s geometry, and the staple design [41]. Furthermore, the scaffold routing is found to influence the way staples are integrated into the scaffold: the integration may happen either through the initial binding followed by a collapse event or in a uniform manner [42].

Here, the mRNA-DNA origami variants D1–D3 featured a simple scaffold routing in which neighboring helices are connected at their ends. In contrast, D4–D5 included two scaffold crossovers shifted towards the center, which could be considered as a ”forth and back” routed design. Assuming that DNA origami folding pathways apply also to (m)RNA-DNA origami, and taking into account the mismatch between the preferred helical pitches of RNA and DNA duplexes, the scaffold routings of D1–D3 might be more suitable for structures adopting 10.5 bp per helical turn. Since the collapse event is global, small structural mismatches might be easier to accommodate, as opposed to the uniform integration required for D4 folding. Interestingly, changing the helical symmetry to 11 bp per turn (D5) helped to overcome any constraints, resulting in the most rigid structure. In addition, when comparing D1 and D2, the staple CO density seems to have a large influence on the folding yield.

Moreover, the CO density has been found to greatly impact the stability of DNA origami structures [31, 43, 44], which is in line with our observations. Despite the similar staple CO density (42 bp vs. 44 bp intervals), D2 was found to be slightly more susceptible to RNase H degradation than D5R-n, most likely caused by the scaffold routing. The lack of the two scaffold COs in the center of the structure may lead to a less compact structure, while simultaneously increasing the flexibility, and by this, the structure becomes more accessible to endonucleases.

The contribution of the helical geometry is reflected in particular upon treatment with DNase I, with D5R-n outperforming D2. Since the activity of DNase I is known to depend on the width of the minor groove [45], D2 is considered more prone to digestion, assuming that the formed mRNA-DNA double helix indeed adopts a helical pitch of 10.5 bp per turn. Indeed, when comparing the electrophoretic mobility of both variants, sharp bands were observed for D5R-n, while the bands for D2 showed a smearing behavior, suggesting degradation of the DNA staple strands. The degradation could also be monitored in the A647 channel, which tracks the A590 fluorophore. A release of A590 was also observed for D5R-n, which could be attributed to the DNase I degradation of the DNA-DNA duplexes formed between the fluorophore-carrying strand and the core staple overhangs.

In general, packaging the mRNA into the presented origami structures was found to make it more stable compared to its free solution form. However, for its applicability, the mRNA must also be readily translated. The translation efficiency was found to depend on (at least) two factors, which are crucial to consider during the design process. First, the translation needs to be initiated, generally through the binding of a preinitiation complex to the 5’-cap, followed by the ribosome recruitment and scanning for the AUG start codon [34]. To this end, the 3’-poly(A)-tail and the 5’-cap region should be placed in close proximity. Additionally, the base-by-base scanning for AUG recognition requires mRNA to be present in a single-stranded form [34]. When integrating the 5’-cap region into the core (D5), no translation was observed when the variant was subjected to the reticulocyte lysate. A similar behaviour has been observed for structures lacking the 5’-cap [24].

Secondly, the translation can only proceed given that the assembled structures unfold. Even though both D2 and D5R-n featured a free, single-stranded 5’-cap region and a poly(A)-tail, the translation efficiency of D2 was found to be on average 1.5*×* higher than that of D5R-n (*P* = 0.045), and *in vitro*, the difference was even more pronounced (*P* = 0.011, C+L, 24 h). We therefore suggest a dependency between the denaturing process and the internal design including the position of the scaffold crossovers and, more importantly, the presence of free scaffold loops. To this end, D2 possessed free scaffold loops at the end of each helix, aiding to release potential strain at the respective ends of the structure and subsequently promote unfolding.

The accessibility of the 5’-cap region is also essential *in vitro*, which in return requires efficient release of the mRNA-DNA origami from its protective coating after the uptake of the complexes into cells. We observed a significant increase in transfection efficiency for mRNA-DNA origami upon complexation with CCMV capsid proteins, a property of plant viruses previously reported for DNA origami [38] and oligonucleotides [46, 47]. The CCMV coating (C) was found to outperform the commercially available transfection agent LP2000. Since the combination C+L showed a similar transfection efficiency, CCMV is suggested to be the driving force for the enhanced uptake.

Recent investigations on the transfection efficiency have demonstrated that not only the mass and the size of the nanoparticle, but also its shape (often referred to as aspect ratio, *i.e.*, the ratio between the long and the short axis of the nanoparticle) [48–51], play a role in uptake. For rod-like DNA origami nanoparticles, an aspect ratio of 1–3 and a length between 50–80 nm were found to be favorable [48]. The presented complexed mRNA-DNA origami variants had, based on the dimensions of the two populations obtained from TEM (Figure 3b), aspect ratios of 1.56 and 2.50 for D2, and 2.25 and 2.84 for D5R-n, and should therefore be efficiently internalized by the cells. Nevertheless, it is notable that for both C and C+L samples, D5R-n showed a slightly higher uptake than D2. This might be attributed to the length of the complexed structures, since locally flat parts at a nanoparticle’s surface may facilitate membrane adhesion at small adhesion strengths [52].

The uptake of nanoparticles commonly follows the endocytotic pathway, however, it is dependent on the cell line as well as on the shape and chemical composition or the biological origin of the nanoparticle [53, 54]. To this end, the uptake can be distinguished between macropinocytosis, clathrin-mediated, and caveolin-mediated endocytosis [6]. Several viruses, including Simian virus 40 [55] and cowpea mosaic virus [56], follow the caveolin-mediated pathway which might be preferential due to the avoidance of the lysosome. In HeLa cells, CCMV is preferentially internalized through the clathrin-mediated endocytosis, though caveolin-mediated endocytosis can occur in case of rod-like structures. During clathrin-mediated endocytosis, the formed vesicle fuses with an early endosome [53].

After internalization, the subsequent release of nucleic acids from the virus capsid shell as well as from the endosome is not well studied. Even the complexation of the plain mRNA with CCMV virus capsid proteins drastically decreased the translation into EGFP when compared to mRNA transfected with LP2000 (Supplementary Figure S21a). A similar behavior of mRNA-loaded virus-like particles has been reported in baby hamster kidney cells (BHK-21), most likely arising from a poor release of mRNA from the CCMV shell [57]. However, by using LP2000 as an additional transfection agent on top of CCMV coating (C+L), we were able to recover ca. 15 % of the EGFP signal (Supplementary Figure S21a).

To further investigate the importance of the disassembly of the capsid shell upon translation, C-samples were incubated with heparin before subjection to reticulocyte lysate. Heparin acts as a negative competing agent, which should therefore trigger the release of mRNA/mRNA-DNA origami from the virus capsid shell (Supplementary Note S11). When complexed structures were treated with an excess of heparin before extracellular translation, ca. 10 % of the EGFP signal could be recovered for the plain mRNA, but neither D2 nor D5R-n showed any fluorescence, suggesting poor or incomplete release from the capsid coat and a (partially) inaccessible 5’-cap region.

*In vitro*, nucleic acids are considered to be released slowly *via* the small pores of the capsid, however, when they are tightly packed, a more rapid release by rupture is conceivable [58]. LP2000, for one, is suggested to interact with the endosome, resulting in destabilization of the endosomal membrane and subsequent gradual release of the cargo into the cytosol of HeLa cells [59]. Our results suggest a synergistic effect between virus capsid proteins and LP2000 for the delivery and translation of mRNA-DNA origami, due to enhancement of transfection while reducing cell toxicity, and promoting endosomal escape, respectively. Once the 5’-cap region is accessible in the cytosol, the mRNA is translated. In comparison to the approaches that complex several mRNA molecules with Lipofectamine into a single particle [60], it is notable that the complexed mRNA-DNA origami encapsulates precisely one mRNA molecule, and therefore the fluorescence signal originating from such nanocarrier is also locally confined.

In conclusion, our results provide a first set of design rules for the successful application of mRNA-DNA origami, highlighting the importance of the accessibility of the start codon for translation initiation and simple scaffold routing to achieve successful translation. Utilizing the DNA origami technique in combination with virus capsid proteins allows to program the size and shape of the nanocarrier template, which in return can be used to modulate the cellular uptake. Additionally, the electrostatic-based complexation enables the assembly of other virus species, and therefore, possible exploitation of the virus’ innate tropism for delivery. Moreover, precise attachment of cargo molecules onto the mRNA-DNA origami structures as well as functionalization of the virus capsid proteins with targeting molecules or ligands could be used [61, 62]. These features could be harnessed in aiding endosomal escape and in achieving highest possible protection and transfection while simultaneously ensuring full release of the mRNA-DNA origami and the accessibility of the start codon. The modularity of the proposed platform would allow for the further development of multipurpose systems for versatile applications in gene delivery and imaging.

## 3 Methods

### 3.1 Folding and purification of mRNA-DNA origami structures

The mRNA-DNA origami variants were folded isothermally in a one-pot reaction. The 996-nt-long EGFP encoding mRNA scaffold, which, according to the manufacturer is capped at its 5’-end to resemble the naturally occurring Cap-1 structure, was purchased from TriLink Bio Technologies (CleanCap EGFP mRNA, L-7601) and the DNA staple strands were obtained from Integrated DNA Technologies. The annealing reaction was performed in an optimized, variant-specific, buffered environment (’folding buffer’, FOB). During the screening, Tris-based as well as HEPES-based buffers were tested, the specific buffer conditions and thermal gradients used are described in Supplementary Note S1,2. For all folding reactions the final concentration of the mRNA was kept constant at 50 nM, and the staple strands (Supplementary Note S12) were used in 10*×* excess (500 nM). Based on the screening, the final FOB containing 1*×*TAE was supplemented with 5 mM MgCl_2_ (D1, D2), 2.5 mM MgCl_2_ (D3, D4), or 5 mM MgCl_2_ and 1 mM NaCl (D5 variants). The mRNA and staples strands were mixed and annealed in a 15 min incubation at 55 °C, followed by a 10 min incubation on ice before storage at 4 °C.

Spin-filtration was used to remove excess staple strands after the annealing. Briefly, 400 µL of 1*×*FOB were added to the spin-filter (100 kDA molecular weight cut-off, MWCO, Amicon) for washing purposes and centrifuged at 14,000g for 5 min. Subsequently, twice the reaction mixture and the 1*×* FOB were added in 1:1 ratio, first 40 µL, followed by 30 µL, and centrifuged at 6,000g for 10 min. Then, the structures were washed 3*×* with 80 µL of 1*×*FOB and twice with 80 µL RNase-free water (6,000g, 10 min). By inverting the filter into a clean tube (1,000g, 2.5 min), the sample was recovered. The concentration of the purified structure was determined from an absorbance measurement at 260 nm using Lambert-Beer’s Law (Note S13).

### 3.2 Agarose gel electrophoresis

Agarose gel electrophoresis was used to monitor the folding of the mRNA-DNA origami as well as the removal of unbound staple strands after purification. To this end, a 3.5 % (w/v) agarose gel (1*×*TAE buffer, 11 mM MgCl_2_) was run at 90 V for 45 min in running buffer containing 1*×*TAE and 11 mM MgCl_2_. This technique was furthermore used to observe the binding interaction between the nucleic acids and the virus capsid proteins as well as to study the intactness of the structures upon nuclease treatment, however, the agarose concentration was decreased to 2 % (w/v). All samples were mixed with 40 % (w/v) sucrose (1:5) or 6*×* gel loading dye (for samples not containing A590), and EtBr (final concentration 0.46 µg mL*^−^*^1^) was used for staining. The nucleic acids were imaged under both ultraviolet light (EtBr) and, if applicible, red light (A647 channel) using a ChemiDoc MP system (Bio-Rad).

### 3.3 Virus coating of mRNA-DNA origami

The complexation of mRNA-DNA origami with CCMV capsid proteins was performed according to ref. [37]. Briefly, CCMV particles were dialysed overnight in Slize-A-Lyzer Mini Dialysis cups (3.5 kDa MWCO, Thermo Scientific) against 50 mM Tris-HCl buffer, pH 7.5 supplemented with 500 mM CaCl_2_ and 1 mM dithiothreitol (DTT), followed by a 6 h centrifugation step (21,000*g*, 4 °C) to pellet the RNA. The supernatant containing the capsid proteins was subjected to another overnight dialysis against ’clean buffer’ (50 mM Tris, pH 7.5 supplemented with 150 mM NaCl and 1 mM DTT). To determine the concentration of the isolated capsid proteins, the absorbance at 280 nm was measured (extinction coefficient 23,590 M*^−^*^1^ cm*^−^*^1^) on a BioTek Eon Microplate Spectrophotometer using a Take3 plate (2 µL sample).

Subsequently, the virus capsid proteins were mixed with purified mRNA-DNA origami in 1*×*FOB in 1:1 ratio resulting in a complexation buffer containing 45 mM Tris, 75.5 mM NaCl, 10 mM acetic acid, 2.5 mM MgCl_2_, 0.5 mM DTT and 0.5 mM EDTA, and incubated for at least 1 h at 4 °C. For the evaluation of the coating by AGE and TEM, the final origami concentration after complexation was 7.5 nM, for the enzymatic stability studies, complexation was performed at 10 nM and for *in vitro* cell assay 31 nM were used.

### 3.4 Enzymatic stability studies

The stability of plain mRNA, as well as plain and complexed mRNA-DNA origami structures in nuclease-rich environment was evaluated using RNase A (from bovine pancreas, Sigma Aldrich, 10109169001), RNase H (NEB, M0297S) and DNase I (from bovine pancreas, Sigma Aldrich, D4263). To this end, 2 µL of the enzyme stock (range 0–500 U mL*^−^*^1^) were added to 15 µL of the sample solution. The final buffer contained 33.75 mM Tris, 7.5 mM acetic acid, 0.375 mM DTT, 0.375 mM EDTA, and after the salt concentration (3 µL, NaCl, CaCl_2_, and MgCl_2_) was adjusted for each sample to ensure optimal conditions for the enzyme, the resulting final sample concentration is 7.5 nM (20 µL). For samples containing plain mRNA, 22.5 nM were chosen as final sample concentrations for easier detection. For both RNase H and DNase I, the NaCl concentration was kept constant at 56.6 mM. Nevertheless, the MgCl_2_ salt concentration was increased to 10 mM for RNase H, and to 5 mM for DNase I. The DNase I buffer was additionally supplemented with 1 mM CaCl_2_. Incubation with RNase A was performed by increasing the NaCl concentration to 500 mM. All enzymes were incubated with the sample at 37 °C for 20 min, after which the sample was placed on ice, mixed with 40 % sucrose (4 µL) and loaded on a 2 % (w/v) agarose gel. Analysis of the band intensities and *R_f_* for RNase A was performed with ImageJ and Origin. *R_f_*was determined as [inline1] , where *r* is the measured distance between the leading band of the treated sample and the gel pocket, while *r*_0_ is the migration distance obtained for the untreated origami sample.

### 3.5 Translation studies using reticulocyte lysate

The capability of different mRNA-DNA origami structure variants to translate into EGFP was studied using a Retic Lysate IVT Kit (Thermo Scientific), following the manufacturer’s instructions with slight modifications. Briefly, 17 µL of reticulocyte lysate (obtained from rabbits) were supplemented with 1 µL amino acid mix (methionine, leucine at concentrations of 1.25 mM), 1.25 µL 20*×* translation mix (1.6 M CH_3_CO_2_K, 10 mM Mg(CH_3_COO)_2_, 200 mM creatine phosphate and 0.5 mM of both methonine and leucine) and 0.25 µL nuclease-free water. Finally, 5 µL of nucleic acid template were added, resulting in total nucleic acid content of 50 ng for mRNA-DNA origami variants or 5– 12 ng for plain mRNA. For reactions studying the strand-displacement, D5R-6 and D5R-10 were incubated for 10 min with 10*×* excess of invader strands. The samples were vortexed gently followed by brief centrifugation before incubating them for 2 h in a 30 °C waterbath. To digest tRNAs in the reaction mixture, 2.5 µL of 1 mg mL*^−^*^1^ RNase A (Sigma Aldrich, 10109169001) were added for 10 min, after which the reaction was stopped by placing the tubes on ice for at least 5 min. The samples were finally stored at -20 °C.

The outcome of the reaction was analysed using native PAGE (4-20% Mini-PROTEAN TGX Precast Protein Gels, Bio-Rad, 4561096). 1 µL of the retic lysate reaction was diluted with 2 µL 1*×*PBS and mixed with 2*×* sample buffer containing 62.5 mM Tris, 40 % (v/v) glycerol and 0.01 % (w/v) bromophenol blue. The gel was run in 25 mM Tris buffer pH 8.3 supplemented with 192 mM glycine at 200 V for 40 min on ice. EGFP was visualized under blue light (A488 channel) with a ChemiDoc MP system (Bio-Rad). The fluorescence intensities were analysed using ImageJ and Origin, the significance was calculated from averaged triplicate samples with a heteroscedastic *t* -test with a two-tailed distribution.

### 3.6 *In vitro* cell assay

For the cell studies, the HeLa cell line was used. It was maintained in complete media consisting of high glucose Dulbecco’s Modified Eagle’s Medium (DMEM, Sigma-Aldrich) supplemented with 5 % fetal bovine serum (FBS, Gibco). Penicillin and streptomycine (final concentrations 100 U mL*^−^*^1^ and 100 µg mL*^−^*^1^, respectively, Gibco) were added to prevent bacterial contamination. The cells were grown in petridishes (Corning) in a humidified 5 % CO_2_ atmosphere at 37 °C. Once the cells reached a confluency of 90 %, they were passaged by incubation with 0.5 % Trypsin-EDTA solution (Sigma-Aldrich) for 5 min to facilitate detachment.

In order to study transfection, the cells (5,000 cells/well) were seeded on 96-well plate (black/clear bottom, TC treated surface, Thermo Scientific) and incubated for 24 hours. Prior to transfection, the cells were washed with 1*×*PBS. Then, the samples were transfected for 4, 16 or 24 hours in Opti-MEM. To allocate sufficient time for the decomplexation of the samples, Opti-MEM was replaced with complete media and the cells were grown for additional 24 h (4 h and 16 h transfection) or over night (24 h transfection). For analysis, the wells were first washed with 100 µL 1*×*PBS to remove residues of the complete media, and then measured in 100 µL 1*×*PBS.

All samples (N, L, C and C+L) were in a first step either complexed with virus capsid proteins (250*×* excess), resulting in a mRNA-DNA origami concentration of 31 nM, or diluted into complexation buffer, matching the concentration. Subsequently, the samples were diluted 1:1 into Opti-MEM, which, for samples treated with Lipofectamine 2000 (Invitrogen) contained 2 % (v/v) (0.7 µL, D2) or 4 % (v/v) (1.4 µL, D5R-n, mRNA) of the transfection agent, at a minimum of 5 min before addition of 20 µL per well, corresponding to 100 ng of RNA.

The success of the transfection was evaluated by measuring the fluorescence emission of both EGFP (520 nm) and A590 (616 nm) in a microplate reader (Cytation 3, BioTek), using excitation wavelengths of 490 nm and 584 nm, respectively. For visualisation, the samples were imaged at 20*×* magnification in bright field (BF) and with EGFP (470 nm) and AF594 (590 nm) filter sets using a Zeiss Axio Observer Z1 microscope. The exposure times for the fluorescence channels was set to 300 ns, and the intensity to 70 % (EGFP) and 65 % (AF594), with exception of plain mRNA samples treated with LP2000 (75 ns exposure, 25 % intensity).

For data treatment, the data from triplicate samples was averaged and normalized by subtracting 90 % of the buffer signal. The significance was calculated with a heteroscedastic *t* -test with a two-tailed distribution.

### 3.7 Atomic force microscopy

A 20 µL droplet of 10 nM origami solution diluted in 1*×*TAE buffer supplemented with 12.5 mM MgCl_2_ was deposited onto a freshly cleaved mica substrate (Electron Microscopy Sciences). The sample was incubated for 1 minute, after which it was washed three times with 100 µL RNase-free water which was immediately plotted away. A steady nitrogen stream was used to dry the sample, after which it was immediately imaged. The measurements were performed in air using ScanAsyst Air mode in combination with ScanAsyst-Air probes (Bruker) on a Dimension Icon AFM (Bruker). The AFM images were processed in NanoScope Analysis v. 1.90, and for easier comparison the height scale was adjusted.

### 3.8 Transmission electron microscopy

Both plain mRNA-DNA origami as well as samples complexed with capsid proteins were prepared at a concentration of 7.5 nM, while for complexed RNA samples 22.5 nM were used. After depositing a sample-containing droplet (3 µL) for 3 min on a plasma cleaned (15 s oxygen plasma flash, NanoClean 1070, Fishione Instruments) Formvar carbon-coated copper grid (FCF400Cu, Electron Microscopy Sciences), the droplet was removed by blotting against filter paper. Subsequently, the grid was negative stained in a two-step procedure using an aqueous 2 % (w/v) uranyl formate solution [27] which had been supplemented with 25 mM NaOH to adjust the pH. After immersing the grid in a 5 µL stain droplet and immediate blotting, the grid was immersed into a 20 µL stain droplet. A final blotting step was performed after 45 s, after which the grids were dried for at least 15 min. The imaging was performed either at an acceleration voltage of 100 kV on a JEOL JEM-2800 electron microscope or at 120 kV on a FEI Tecnai 12 Bio-Twin microscope.

**Supplementary information.** A collection of Supplementary Notes (Note 1–13) which include Supplementary Figures (S1-S23) and Supplementary Tables (S1-S6) can be found in the accompanying .pdf file

## Acknowledgments

The authors acknowledge financial support from the European Research Council (ERC) and ERA Chair MATTER under European Union’s Horizon 2020 research and innovation programme (grant agreement no. 101002258), Emil Aaltonen Foundation and Jane and Aatos Erkko Foundation. We thank Ahmed Shaukat for his assistance with cell culturing. This work was carried out under the Academy of Finland Centers of Excellence Program (2022-2029) in Life-Inspired Hybrid Materials (LIBER), project number (346110). We acknowledge the provision of facilities and technical support by Aalto University Bioeconomy Facilities, OtaNanoNanomicroscopy Center (Aalto-NMC) and Micronova Nanofabrication Center.

## Data availability

All the generated and analysed data of this study are included in this published article or in the supplemented files, and are also available from the corrresponding author upon request.

## Authors’ contributions

Project conceptualization: I.S., S.S., M.A.K. Origami design: I.S., V.L. Characterization: I.S., S.S. Translation using reticulocyte lysate: I.S. *In vitro* cell studies: J.W., S.S., I.S. CCMV production: J.J.L.M.C. Manuscript writing: I.S., V.L. and M.A.K. with inputs from all authors.

## Competing interests

The authors declare no competing interests.

## Notes

### Competing Interest Statement

The authors have declared no competing interest.

