## Supplementary Information for "Folding of mRNA-DNA origami for controlled translation and viral vector packaging"

### Contents


**Page**

Note S1: Optimization of the folding conditions using Tris -based buffers (Supplementary Figures 1–6) . . . S3
Note S2: Folding of D2–D5 using HEPES -based buffers (Supplementary Figures 7–10) . . . S9
Note S3: Rerouting D5 (Supplementary Figure 11) . . . S13
Note S4: Retic lysate assay (Supplementary Figure 12) . . . S14
Note S5: Coating of D2 and D5R-n with CCMV capsid proteins (Supplementary Figure 13) . . . S15
Note S6: DNase I digestion of D5R-n and D2 (Supplementary Figure 14) . . . S16
Note S7: *In vitro* cell studies (Supplementary Figures 15–18) . . . S17
Note S8: Complexation with Lipofectamine 2000 (Supplementary Figure 19) . . . S21
Note S9: Delivery of plain mRNA *in vitro* (Supplementary Figures 20–21) . . . S22
Note S10: MTT assay (Supplementary Figure 22) . . . S24
Note S11: Disassembly of complexed structures (Supplementary Figure 23) . . . S25
Note S12: Staple sequences for D1–D5 (Supplementary Tables 1–5) . . . S26
Note S13: Estimation of DNA origami concentration (Supplementary Table 6) . . . S30

### Note S1: Optimization of the folding conditions using Tris-based buffers

The five mRNA-DNA origami variants, differing in their internal design, were folded using both 0.5×TE buffer and 1×TAE buffer. To screen the negative charge originating from the backbone of the nucleic acids, the buffers were supplemented with NaCl and $MgCl_2$, respectively. For each buffer condition, a specific temperature gradient (heating to 65 °C for 30 s followed by cooling from 65 °C to 30 °C with a ramp of -0.1 °C per 1.5 min and a ramp of -0.1 °C per 2 min between 30 °C to 20 °C) (1) or isothermal folding at 55 °C was chosen to anneal the staples to the structures. Figures S1–S5 supplement the folding of the structures presented in Figure 2.

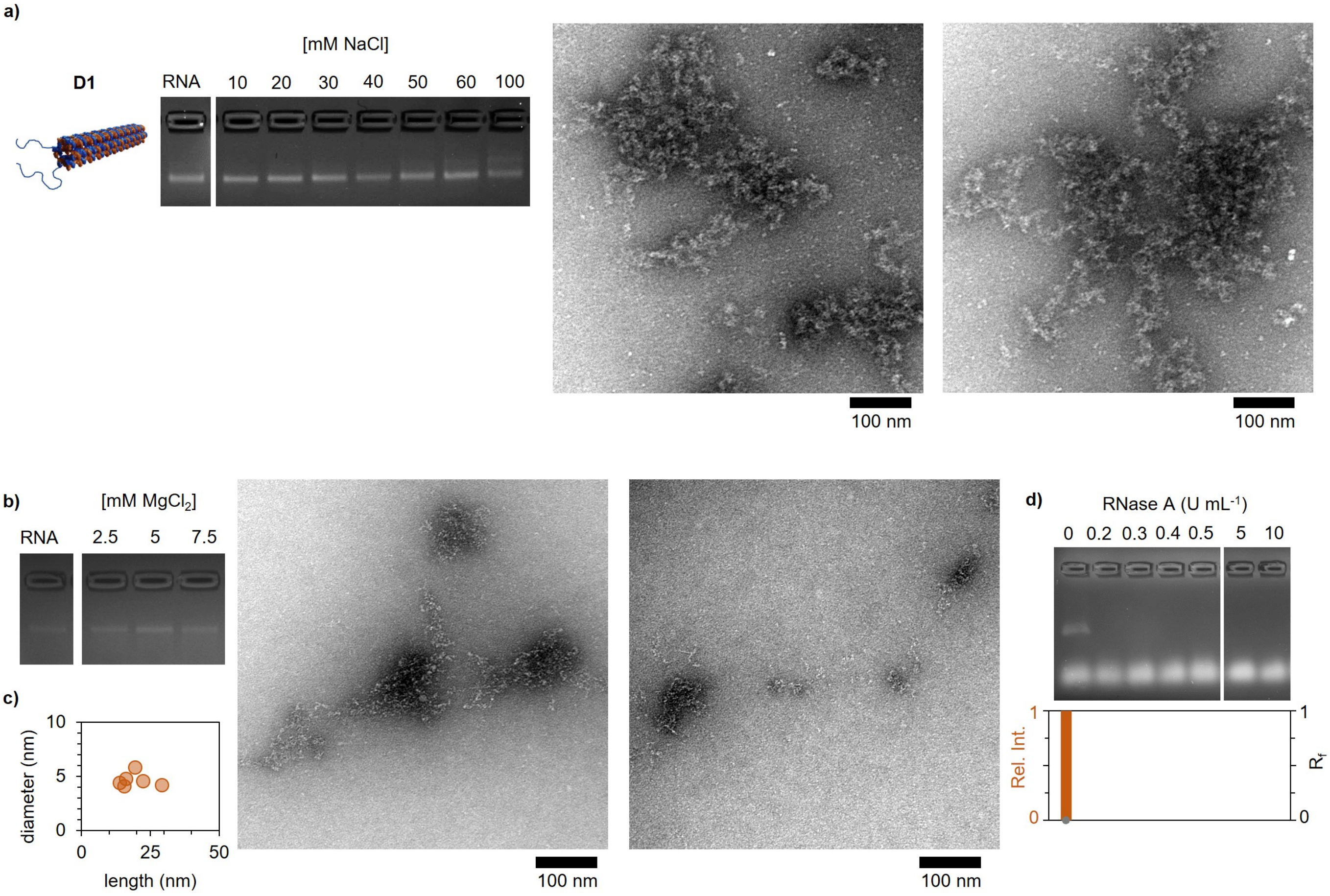


**Figure S1.** The optimal buffer conditions for the folding of D1 were screened **a**, using 0.5×TE supplemented with 0–100 mM NaCl (left), which mainly resulted in unfolded or aggregated structures (right), and **b**, 1×TAE buffer supplemented with 0–7.5 mM $MgCl_2$ (left). Despite the majority of the sample being unfolded or aggregated supplementing with 5 mM $MgCl_2$ (right), a few partially folded structures were observed, **c**, ranging from 13–29 nm. **d**, The low folding yield could also be demonstrated based on treatment of the sample with RNase A (top). The leading band vanishes already when exposed to 0.2 U $mL^{-1}$, so neither the intensity nor the electrophoretic mobility can be determined (bottom).

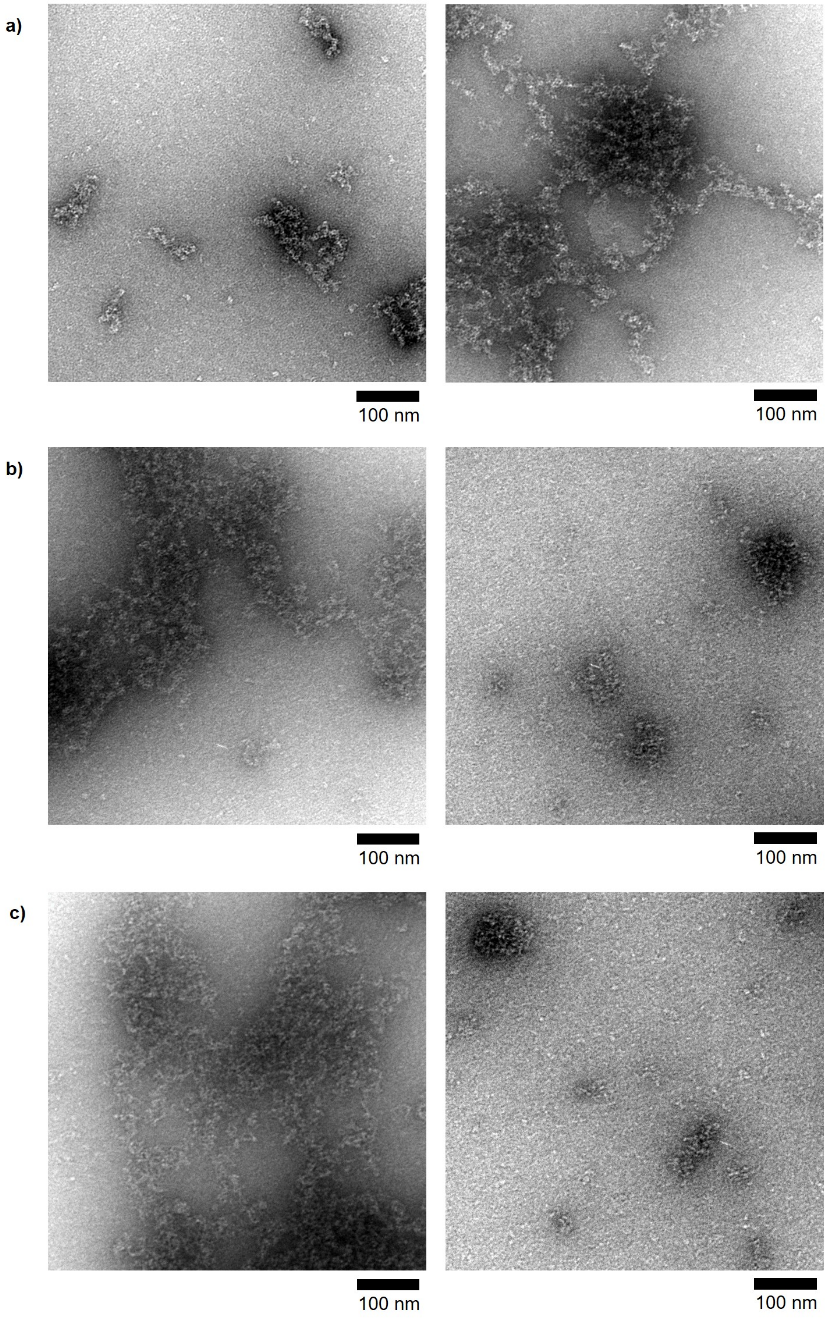


**Figure S2.** Supplementary negative-stain TEM images showing the folding in 0.5×TE supplemented with 30 mM NaCl for **a**, D2, **b**, D3, and **c**, D4.

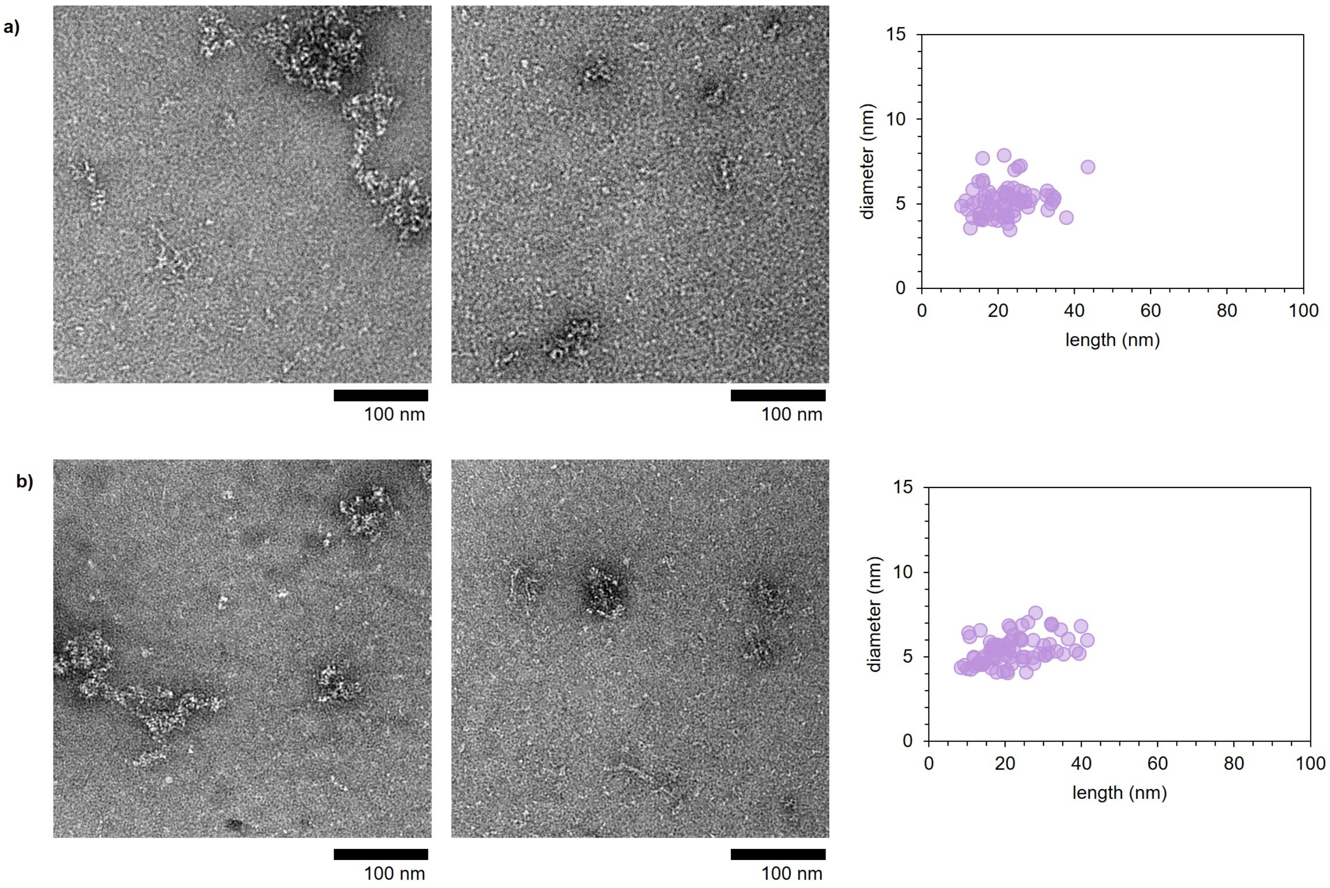


**Figure S3.** Folding of D5 in 0.5×TE supplemented with **a**, 30 mM NaCl and **b**, 100 mM NaCl, characterized with negative-stain TEM (left) and the measured dimensions of the structures (right).

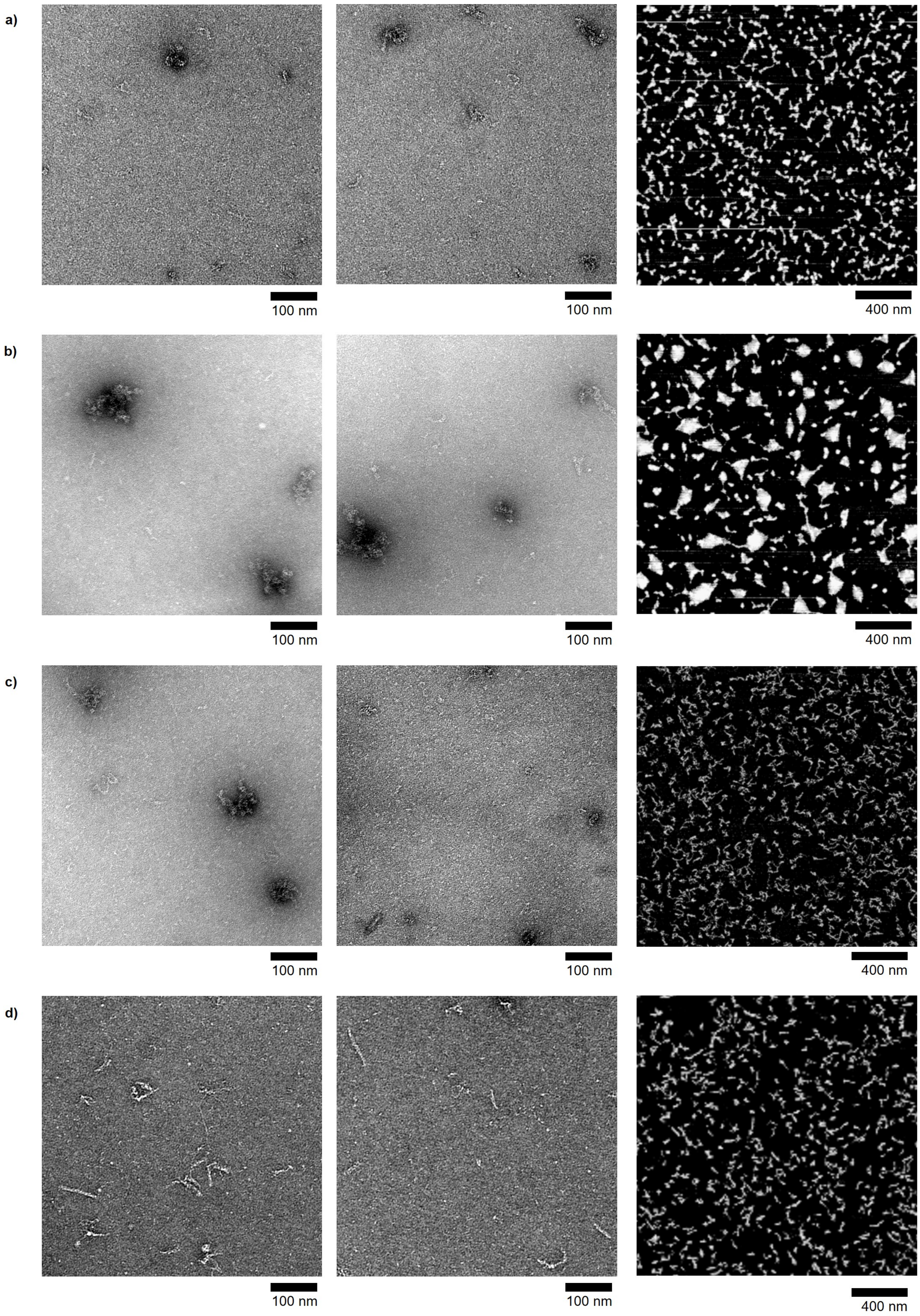


**Figure S4.** Supplementary negative-stain TEM images (left) and AFM (right) showing the folding in 1×TAE supplemented with $MgCl_2$ for **a**, D2 (5 mM $MgCl_2$), **b**, D3 (2.5 mM $MgCl_2$), **c**, D4 (2.5 mM $MgCl_2$), and **d**, D5 (5 mM $MgCl_2$).

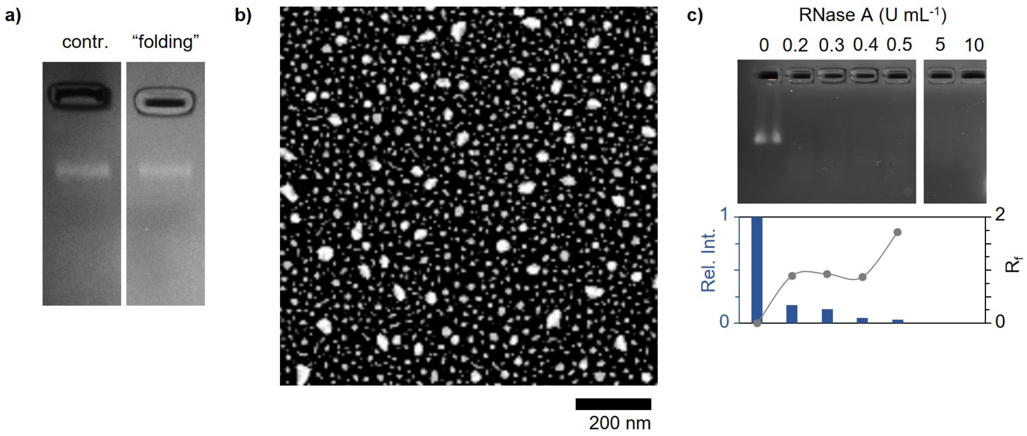


**Figure S5.** The mRNA was subjected to the folding conditions (55 ℃, 15 min in 1×TAE supplemented with 5 mM $MgCl_2$) and analyzed using **a**, agarose gel electrophoresis (AGE) and **b**, AFM. **c**, The stability of the mRNA when exposed to RNase A (top) was evaluated by following the electrophoretic mobility ($R_f$, grey) and the intensity (bar) of the leading band. Note that mRNA was used at three times higher concentration than for mRNA-DNA origami variants (Figure 2i-l).

In order to reduce the formation of dimers, supplementation of the folding buffer (1×TAE, 5 mM $MgCl_2$) with NaCl is tested (Figure S6a), arriving at a final buffer composition of 1×TAE, 5 mM $MgCl_2$, 1 mM NaCl (Figure S6b,c).

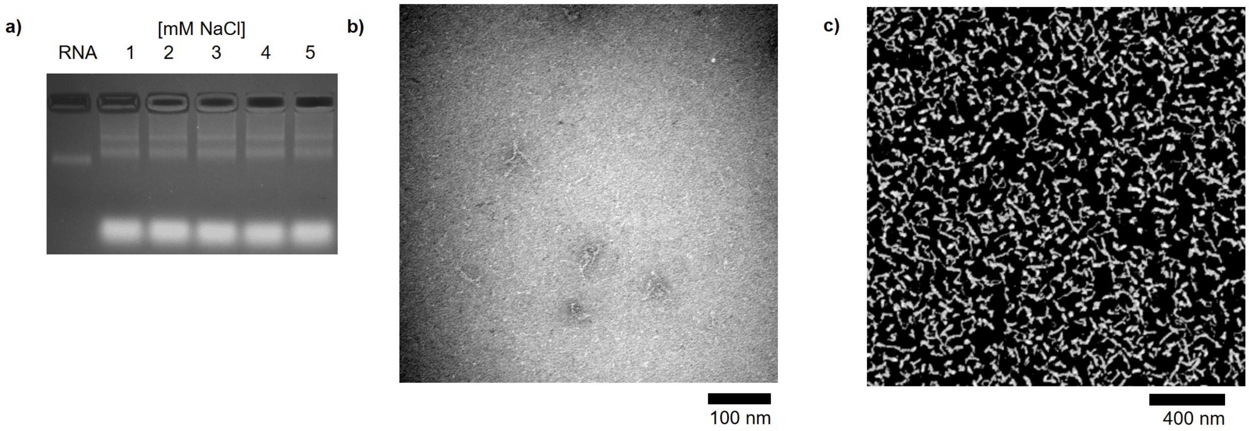


**Figure S6. a,** Optimization of the folding buffer by supplementing with NaCl. D5 folded in 1×TAE, 5 mM $MgCl_2$, 1 mM NaCl analysed with **b,** TEM and **c,** AFM.

### Note S2: Folding of D2–D5 using HEPES-based buffers

All here tested folding conditions are based on previously described protocols. While most protocols are tris-based (1–3), HEPES (10 mM) offers, in combination with KCl for screening, a promising alternative minimizing RNA instability upon high temperatures and divalent cations (4). To investigate the impact of the temperature on the folding process of D2–D5, we tested a temperature gradient by heating the samples to 90 °C for 45 s, followed by cooling from 85 °C to 70 °C with a rate of 45 s °C$^{-1}$, from 70 °C to 29 °C with a rate of 15 min °C$^{-1}$ and from 29 °C to 25 °C with a rate of 10 s °C$^{-1}$ (4) *vs.* isothermal folding for 15 min at 55 °C. As soon as KCl is added into the folding buffer, all designs (Figures S7–10) display a shift in electrophoretic mobility which does not differ significantly between structures folded with (a) the temperature gradient and (b) isothermally. Furthermore, the structures show a tendency for dimerization, as displayed as a band with lower mobility. While persistent for D2 and D5, the dimerization decreases for D3 and D4 at high KCl concentrations. Analysis of the samples (supplemented with 300 mM KCl for D2–D4, and 200 mM KCl for D5) using TEM reveals indeed folding, however, especially D3 and D4 show a tendency for aggregation when folded using the temperature gradient.

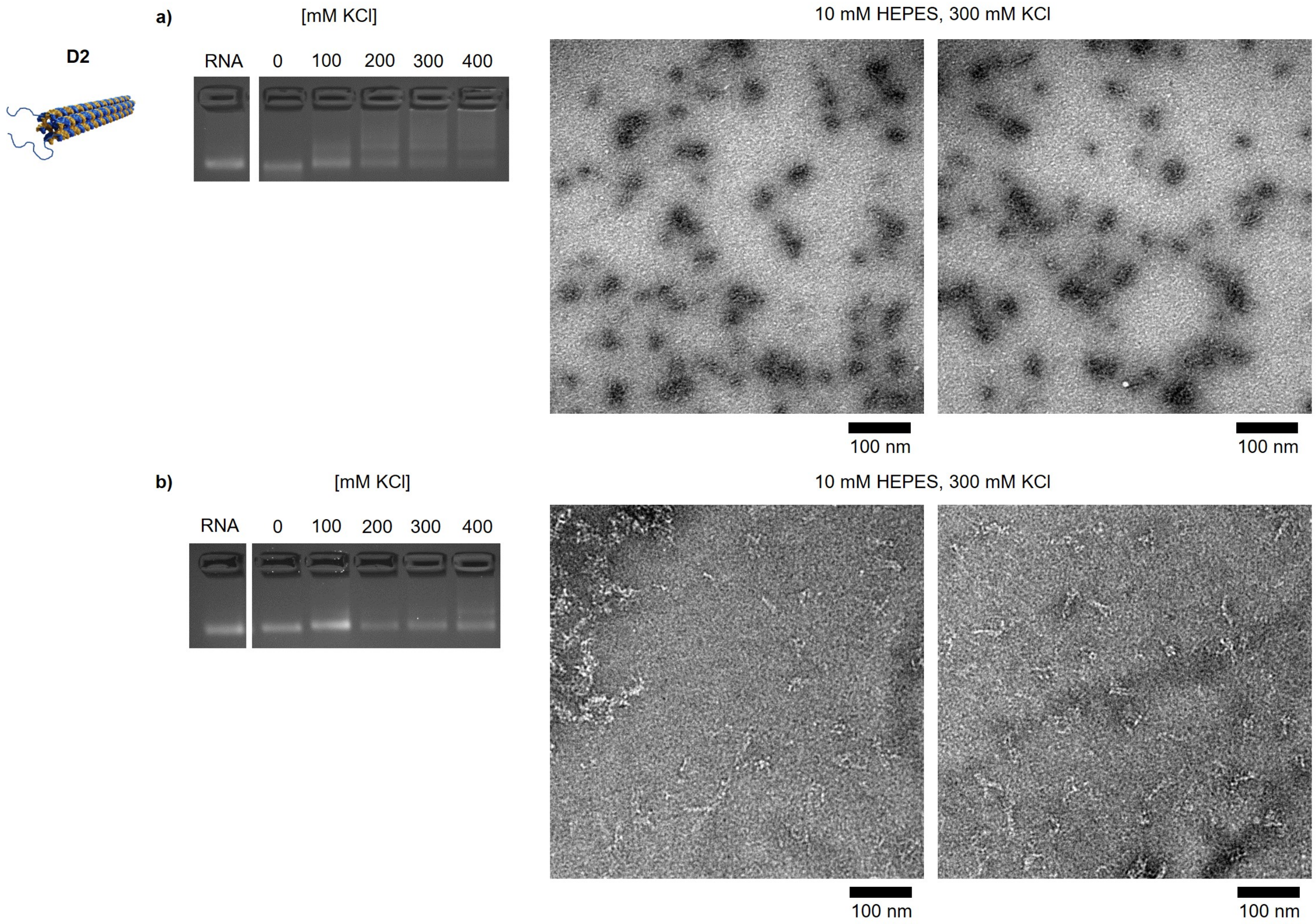


**Figure S7.** KCl screening to supplement HEPES buffer facilitating the folding of D2 when using **a**, a temperature gradient and **b**, isothermal temperature.

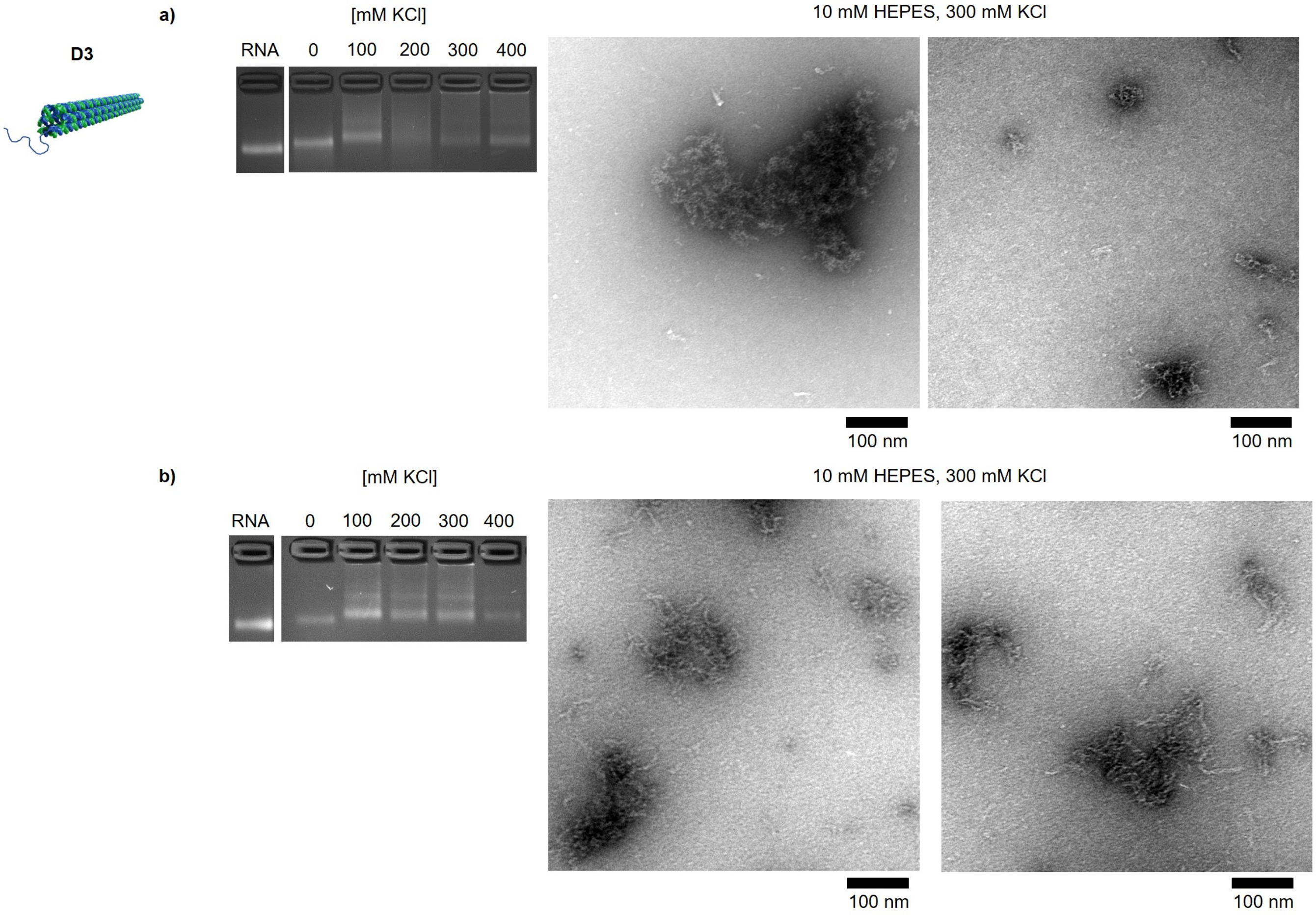


**Figure S8.** KCl screening to supplement HEPES buffer facilitating the folding of D3 when using **a**, a temperature gradient and **b**, isothermal temperature.

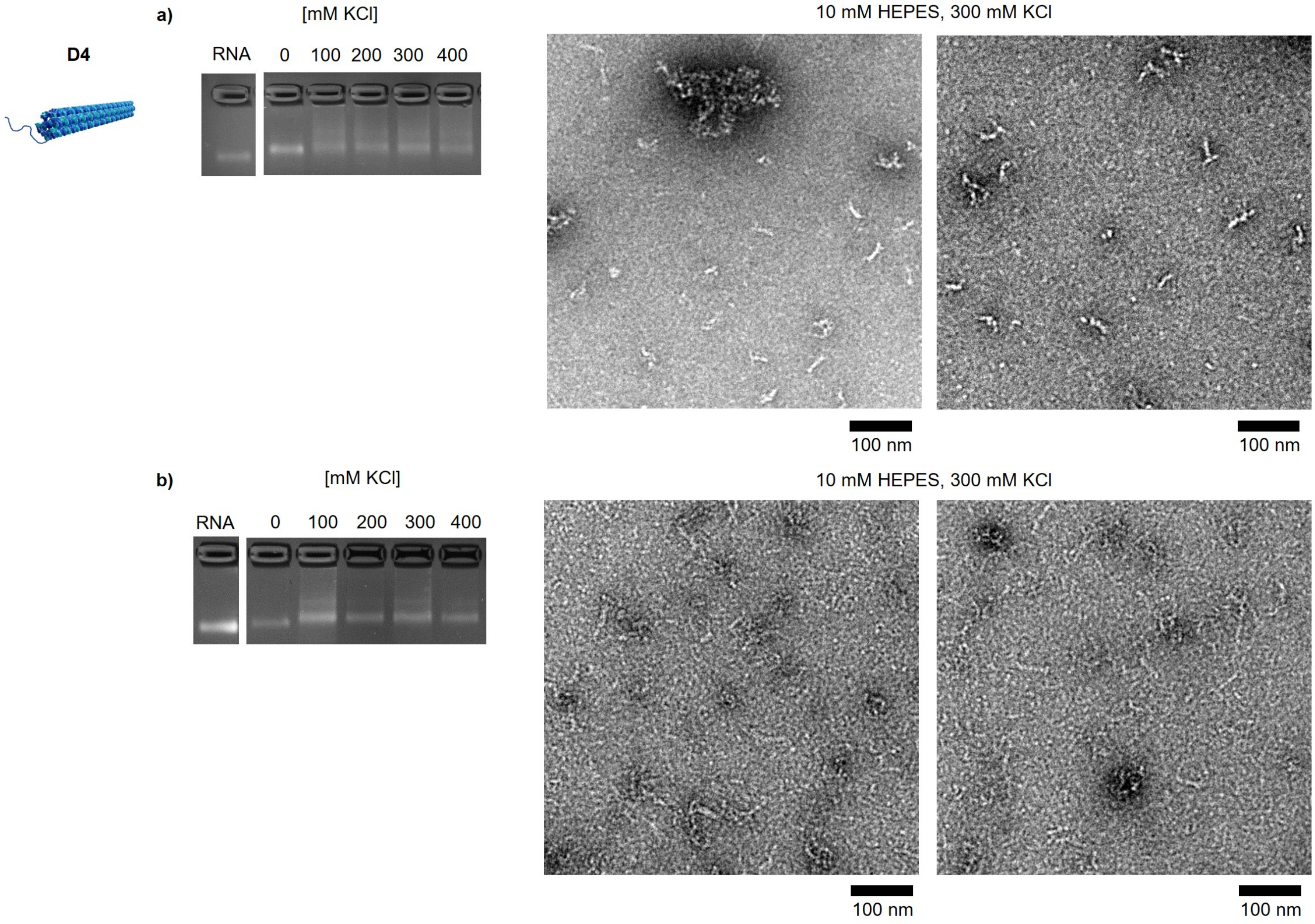


**Figure S9.** KCl screening to supplement HEPES buffer facilitating the folding of D4 when using **a**, a temperature gradient and **b**, isothermal temperature.

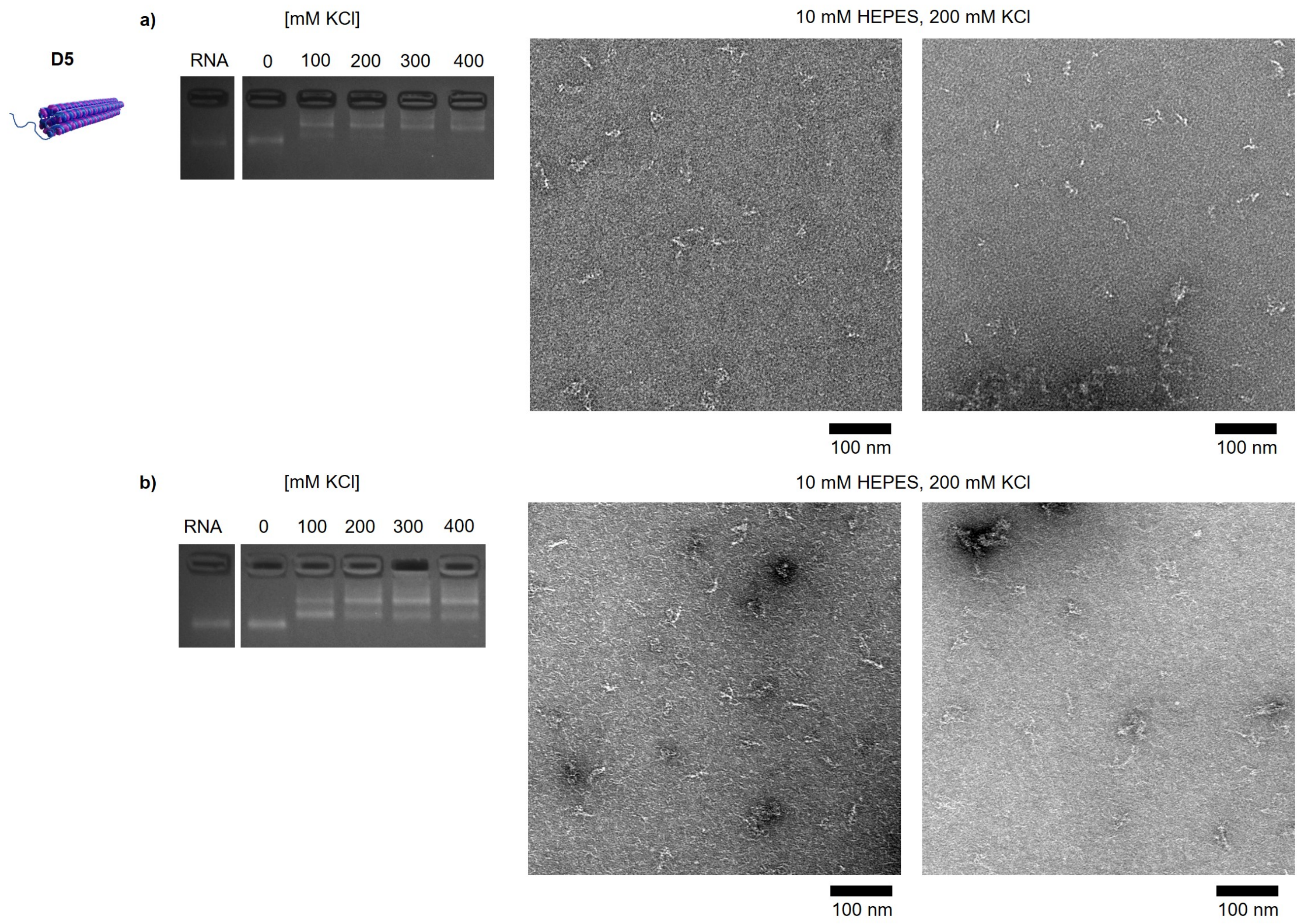


**Figure S10.** KCl screening to supplement HEPES buffer facilitating the folding of D5 when using **a**, a temperature gradient and **b**, isothermal temperature.

### Note S3: Rerouting D5

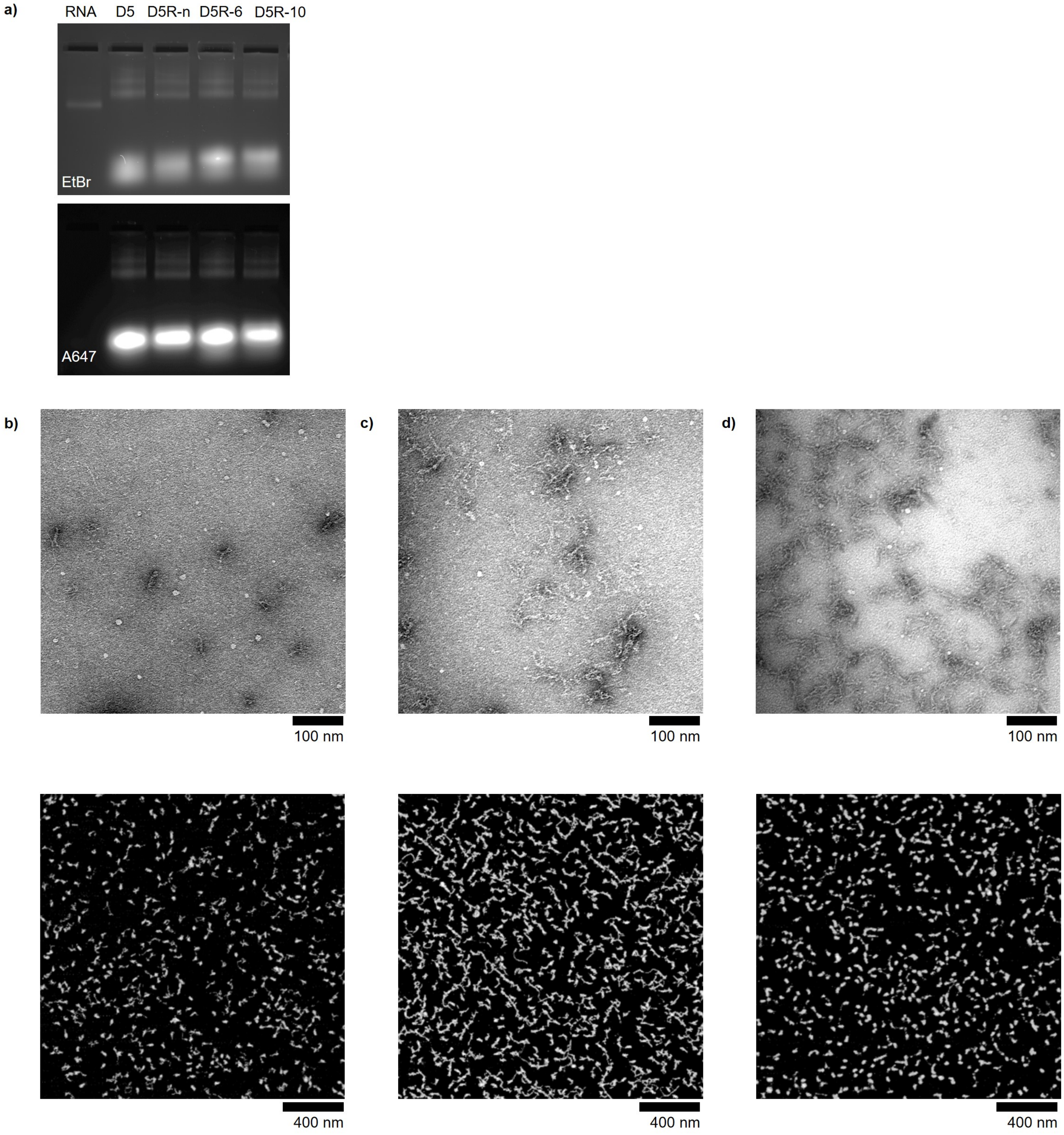


**Figure S11.** The folding of the rerouted structures was evaluated using **a**, AGE as well as TEM (top) and AFM (bottom) for **b**, D5R-n, **c**, D5R-6, and **d**, D5R-10.

### Note S4: Retic lysate assay

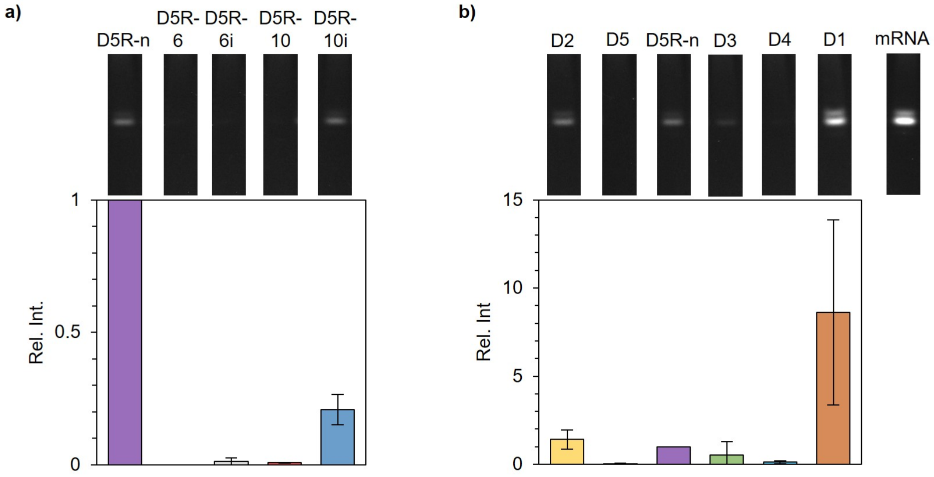


**Figure S12.** Native PAGE was used to monitor the translation of **a**, the rerouted structures D5R-n (purple), D5R-6 (light red), D5R-10 (red). Additionally, the respective invader strands were incubated with the origami variants 10 min before addition to reticulocyte lysate, D5R-6i (grey) and D5R-10i (blue). **b**, GFP signal of unpurified origami structures. All intensities are relative to the intensities of D5R-n.

### Note S5: Coating of D2 and D5R-n with CCMV capsid proteins

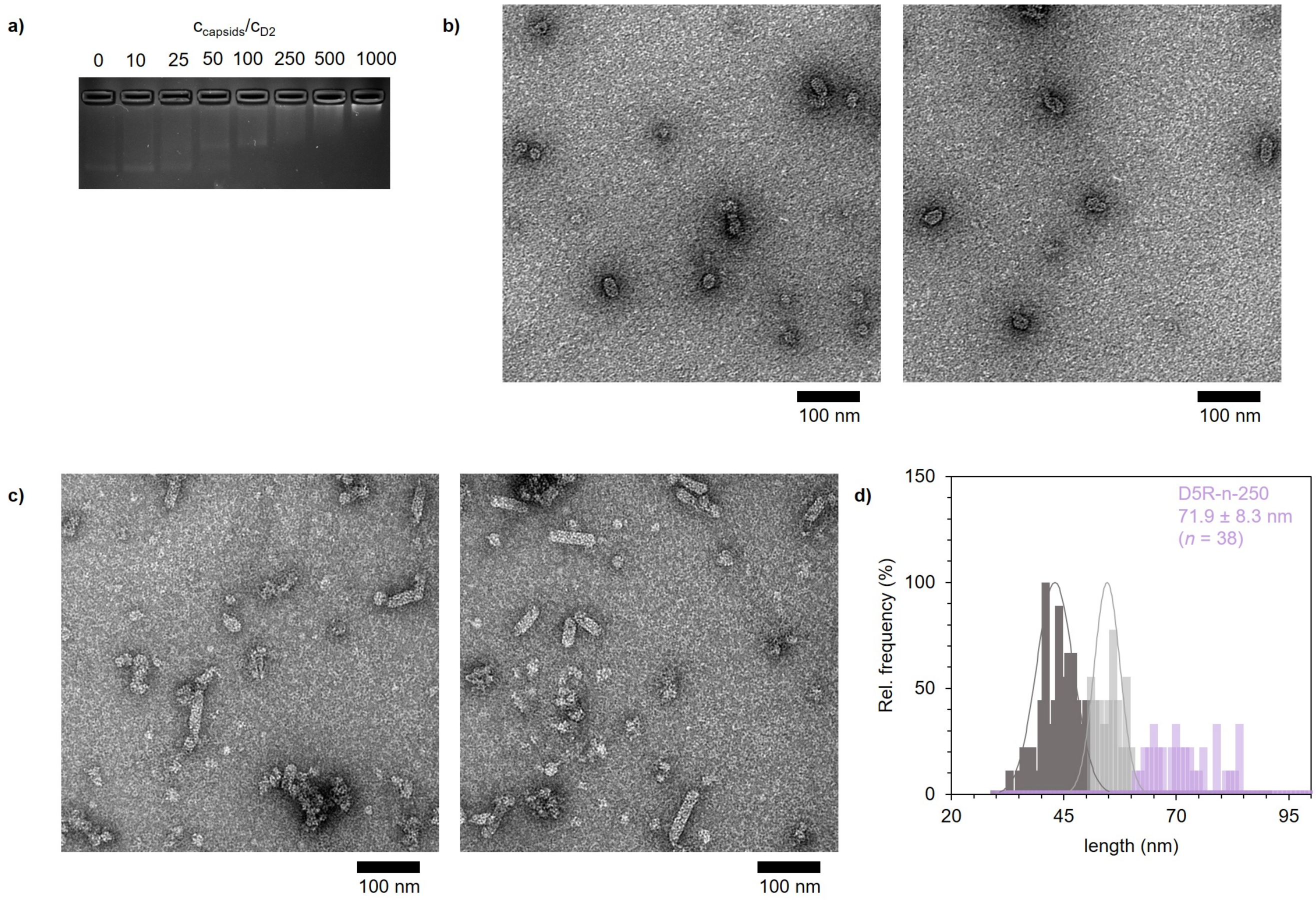


**Figure S13. a**, Complexation of D2 with increasing excess of CCMV virus capsid proteins, resulting in **b**, coated structures (protein excess of 250). **c**, D5R-n complexed with CCMV virus capsid protein excess of 250 shows **d**, a broad size distribution (Figure 4d), including dimer structures (light purple) with an average length of 71.9 nm.

### Note S6: DNase I digestion of D2 and D5R-n

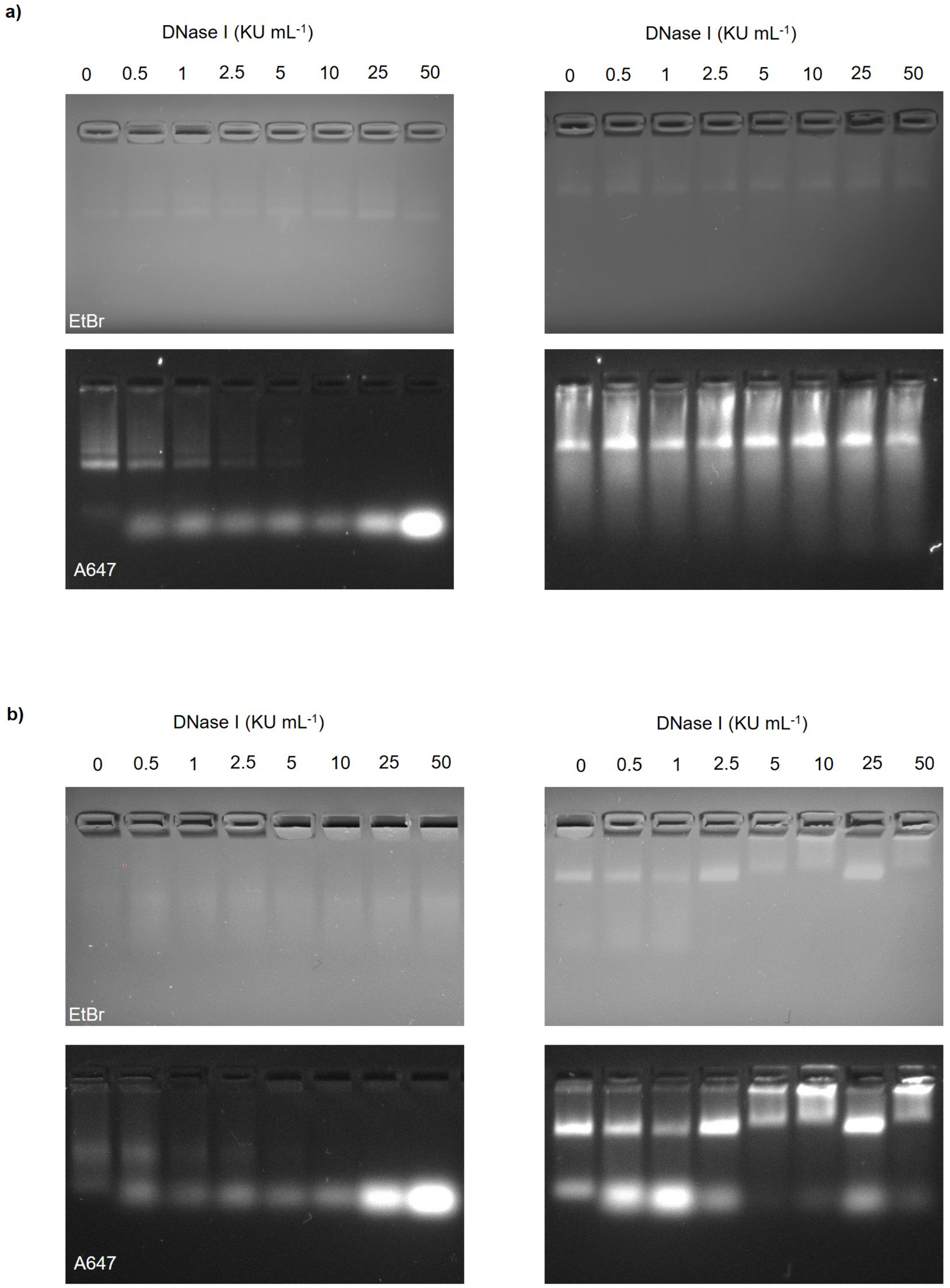


**Figure S14.** The susceptibility of **a,** D5R-n and **b,** D2 against DNase I was monitored by AGE. While plain mRNA-DNA origami variants (left), especially D2 shows degradation, the origami structures were protected when complexed with virus capsid proteins (right).

### Note S7: *In vitro* cell studies

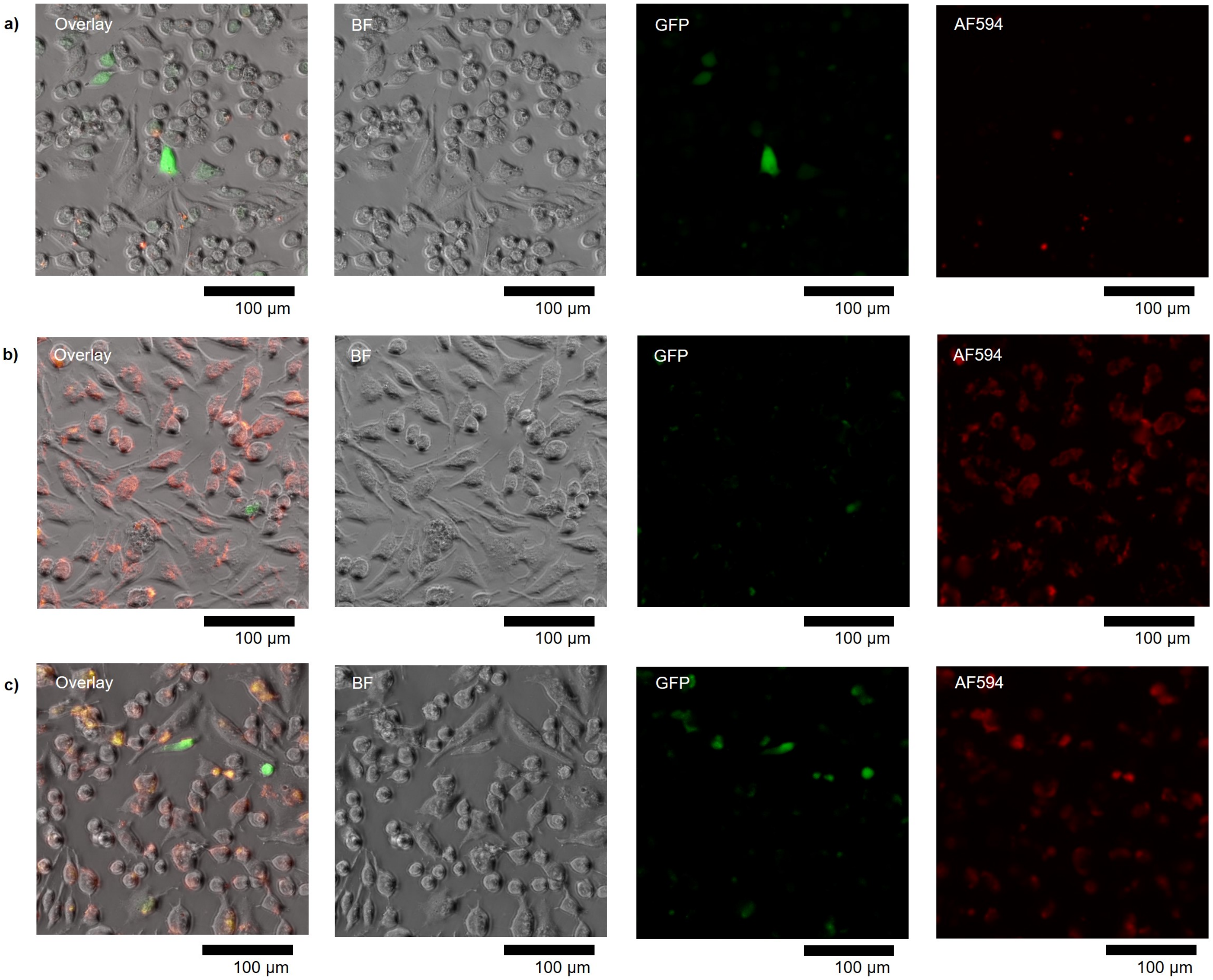


**Figure S15.** Uptake and translation of D2 when transfected for 16 h with **a**, Lipofectamine 2000 (LP2000), **b**, CCMV capsid proteins (excess of 250), and **c**, a combination of capsid proteins and LP2000, showing bright field (BF), GFP and A590 (AF594) channels.

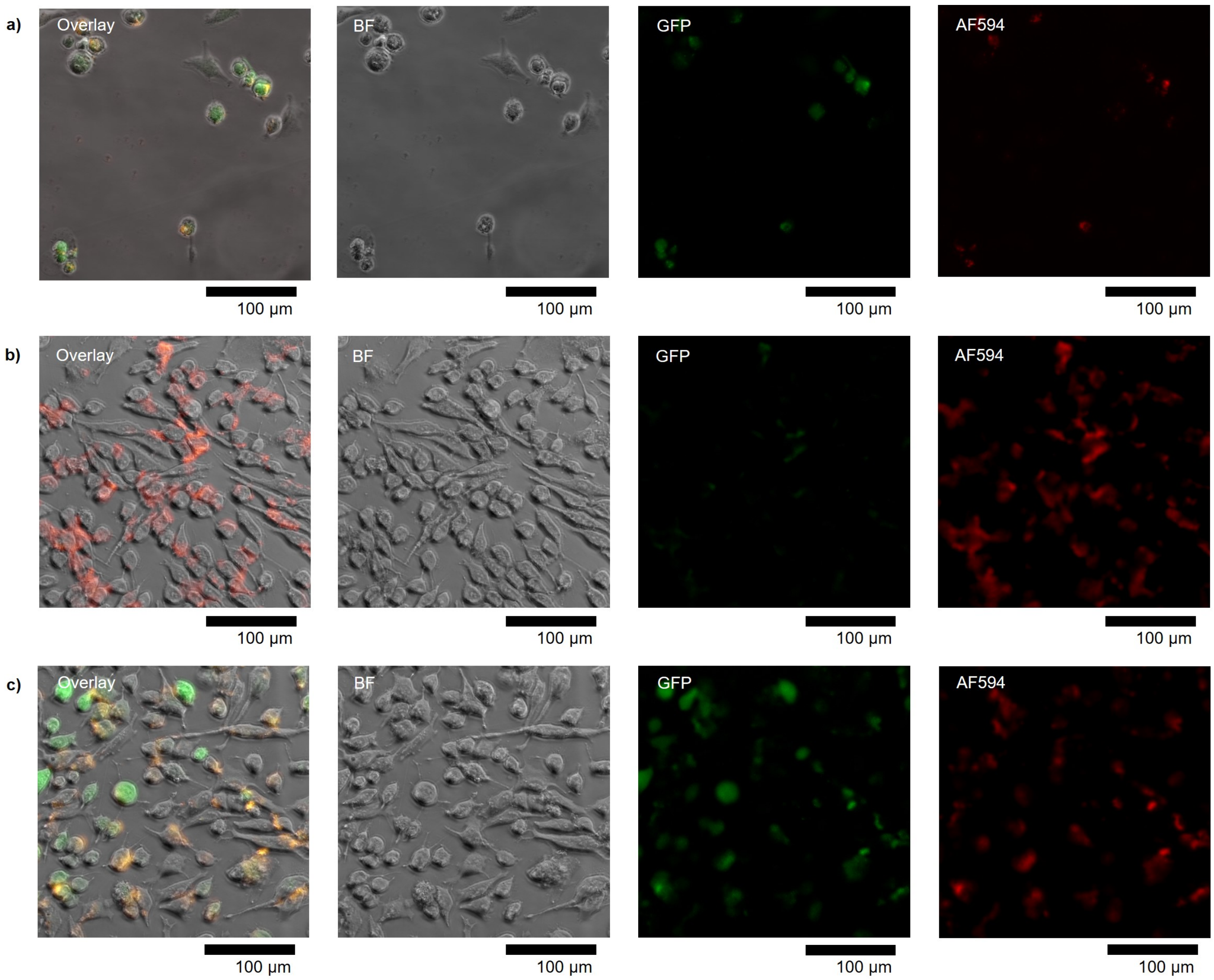


**Figure S16.** Uptake and translation of D2 when transfected for 24 h with **a**, LP2000, **b**, CCMV capsid proteins (excess of 250), and **c**, a combination of capsid proteins and LP2000, showing BF, GFP and A590 (AF594) channels.

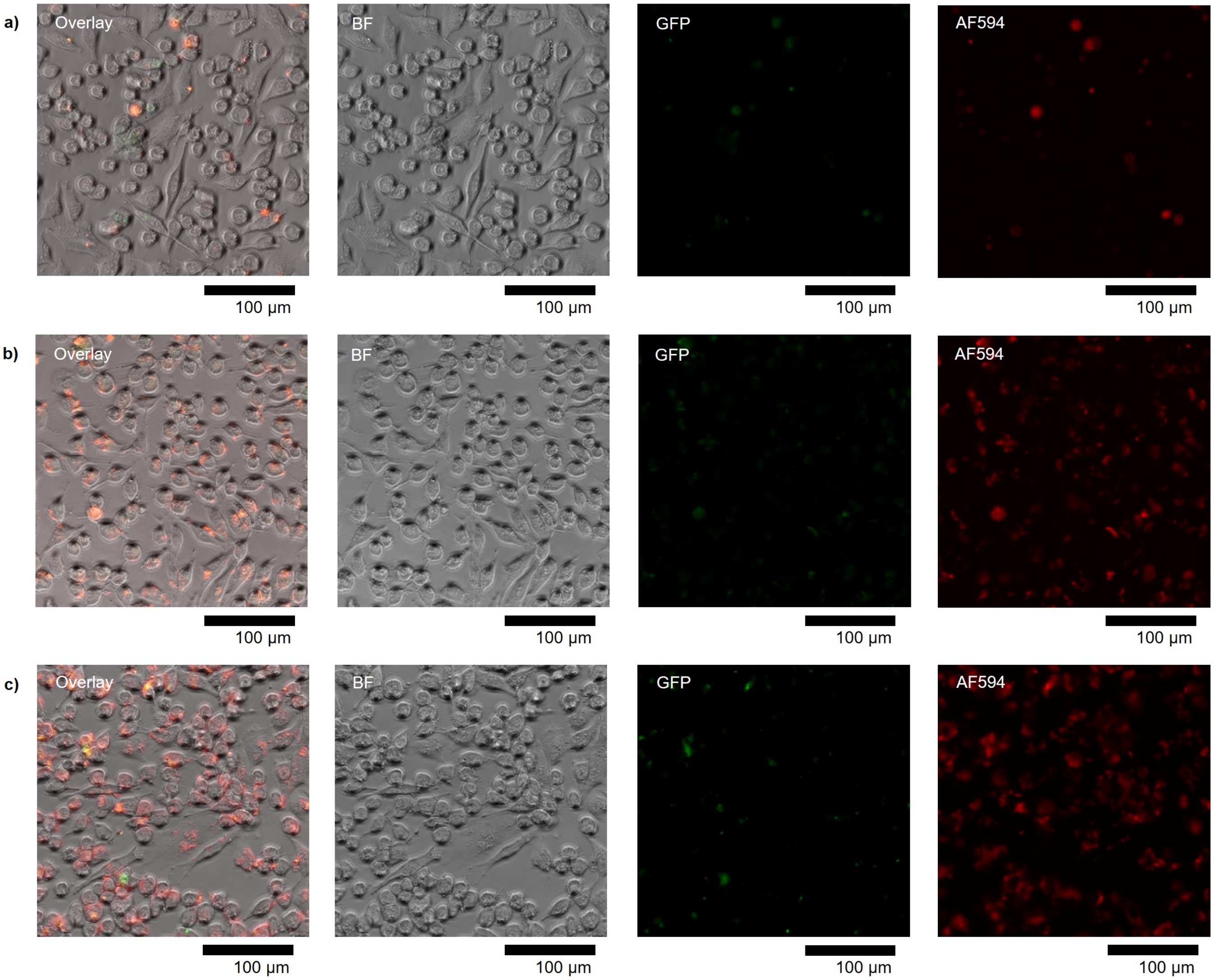


**Figure S17.** Uptake and translation of D5R-n when transfected for 16 h with **a**, LP2000, **b**, CCMV capsid proteins (excess of 250), and **c**, a combination of capsid proteins and LP2000, showing BF, GFP and A590 (AF594) channels.

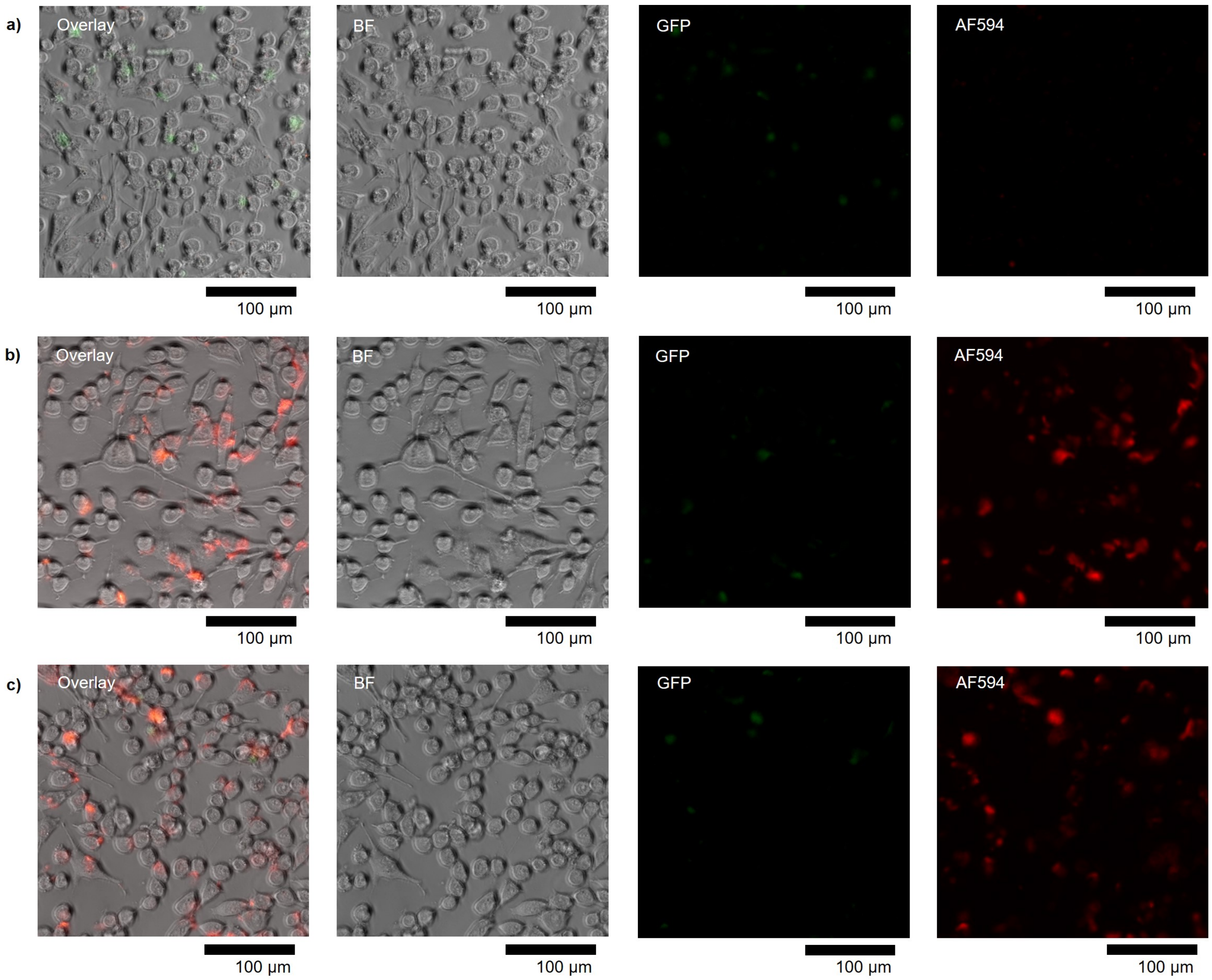


**Figure S18.** Uptake and translation of D5R-n when transfected for 24 h with **a**, LP2000, **b**, CCMV capsid proteins (excess of 250), and **c**, a combination of capsid proteins and LP2000, showing BF, GFP and A590 (AF594) channels.

### Note S8: Complexation with Lipofectamine 2000

For the *in vitro* cell studies, mRNA-DNA origami was characterized with all transfection agents, *i.e.*, not only the complexation with virus capsid proteins (C), but also when incubated with LP2000 (L, Figure S19a) or with both transfection agents (C+L). For the complexation, D2 or D2-250 was incubated for 5 min with LP2000 in Opti-MEM (% v/v), followed by AGE or deposition on a TEM grid for analysis. Increasing the amount of LP2000 above 4 % (v/v) results in aggregation of D2, for the *in vitro* studies 2 % (v/v) was used for D2 and 4 % (v/v) for D5R-n and mRNA. D2-L was found to mainly result in round particles, whereas D2-C+L maintains the virus capsid coat and subsequently its rod-like nature.

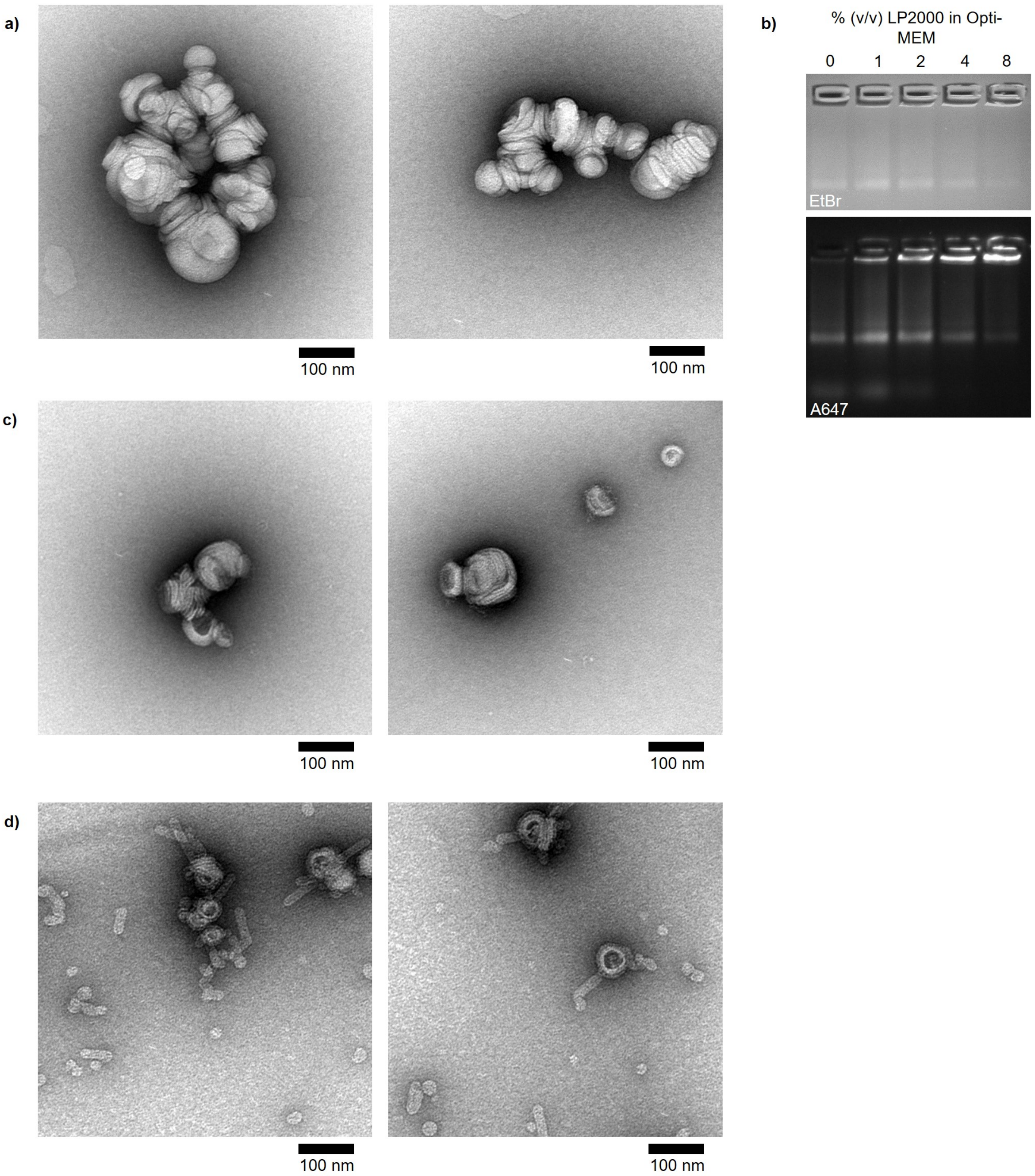


**Figure S19. a,** Negative stain TEM 2 % (v/v) LP2000 in Opti-MEM. **b,** The shift in electrophoretic mobility upon complexation of D2 with LP2000 in Opti-MEM. Negative stain TEM of **c,** D2 complexed with LP2000 and **d**, D2 complexed first with a protein excess of 250, followed by 2 % (v/v) LP2000 in Opti-MEM.

### Note S9: Delivery of plain mRNA *in vitro*

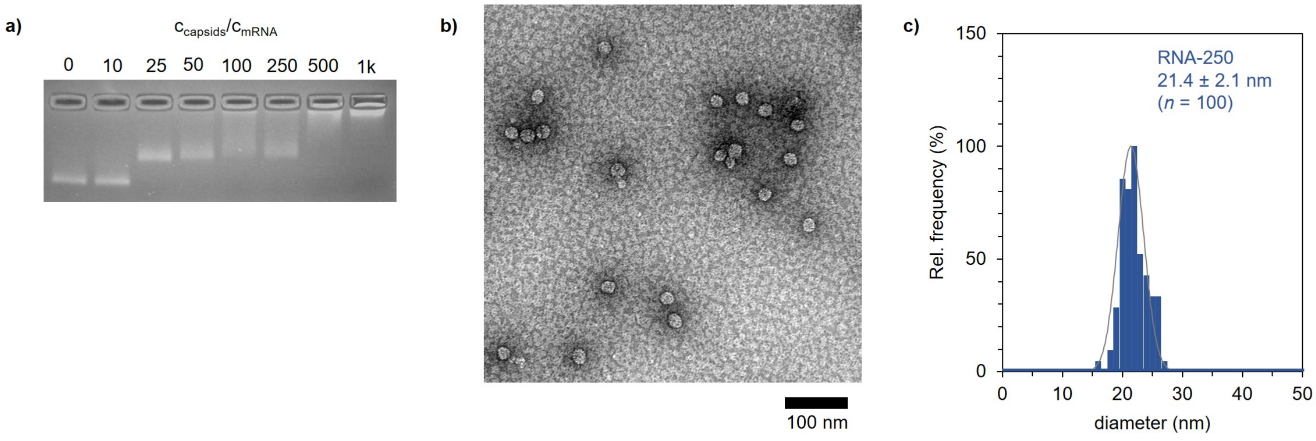


**Figure S20. a,** mRNA complexation with virus capsid proteins results in a shift in electrophoretic mobility. The complexed structures (mRNA-250) appear spherical under **b,** TEM, **c,** with an average diameter of 21.4 $\pm$ 2.1 nm.

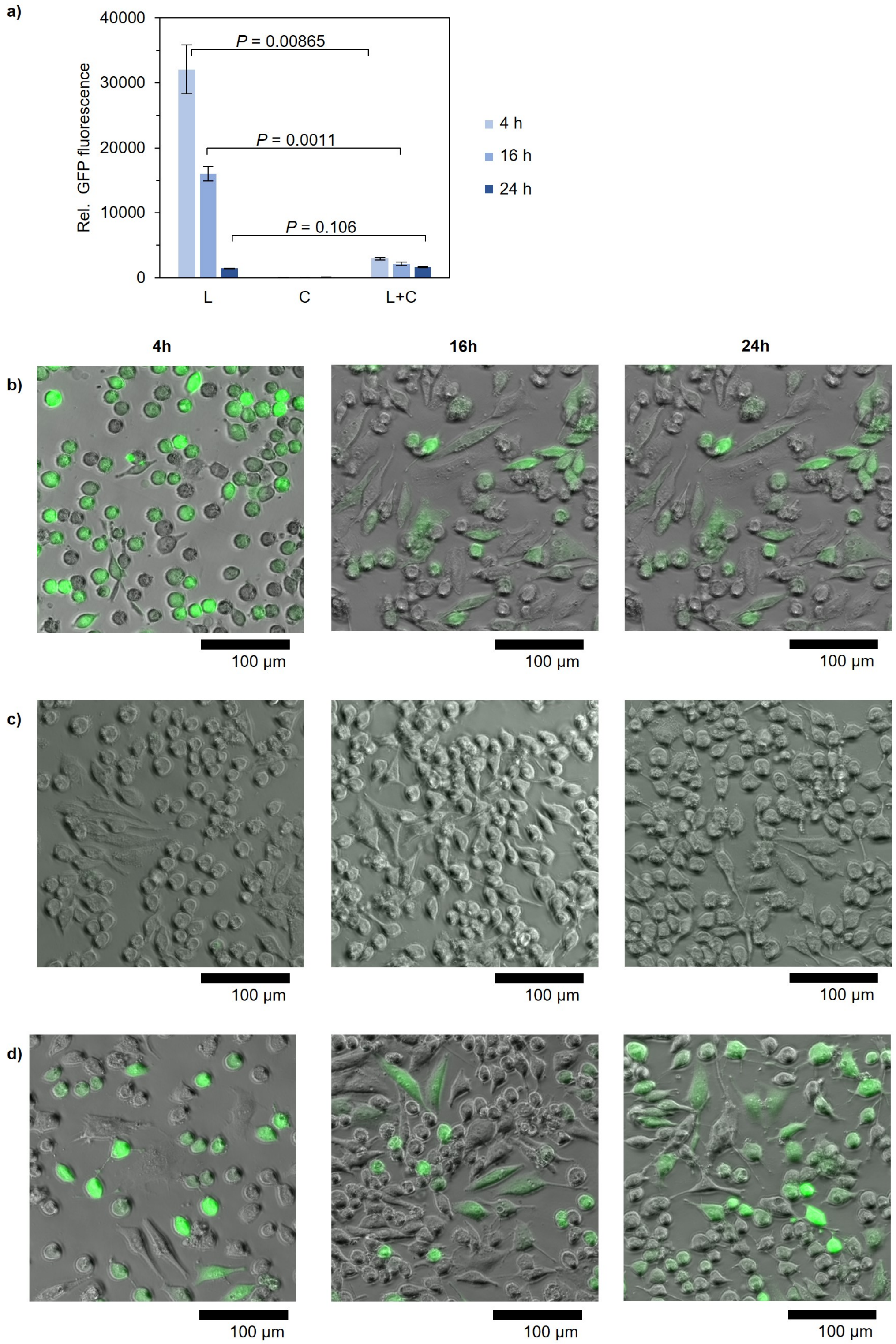


**Figure S21.** The translation of mRNA in HeLa cells was monitored by EGFP's fluorescence signal using **a,** a microplate reader and by fluorescence microscopy for **b,** mRNA-L, **c,** mRNA-C, and **d,** mRNA-C+L when incubated for 4 h (left), 16 h (middle) and 24 h (right).

### Note S10: MTT assay

The cell viability was evaluated using a 3-(4,5-dimethylthiazol-2-yl)-2,5-diphenyl-tetrazolium bromide (MTT) dye assay. 5,000 cells per well were seeded into a 96-well plate 24 h prior to the addition of the samples and incubated at 37 °C in a humidified environment supplemented with 5 % $CO_2$. The media was exchanged to Opti-MEM and the respective mRNA/mRNA-DNA origami samples were added, corresponding to 100 ng of mRNA per well. The incubation was continued for 24 h, after which the media was replaced with 100 μL complete media and the cells were further incubated overnight (corresponding to the 24 h conditions used for transfection studies). An MTT stock of 5 mg $mL^{-1}$ (1×PBS) was diluted into complete media to a final MTT concentration of 0.5 mg $mL^{-1}$. The overnight cell media was subsequently replaced with 100 μL of the prepared MTT containing media and the cells were incubated for 4 h. Then, the media was replaced with 100 μL DMSO to dissolve the formazan crystals prior to analysis with a microplate reader (Cytation 3, Biotek). The absorbance of the triplicate samples was measured at the wavelength of 570 nm.

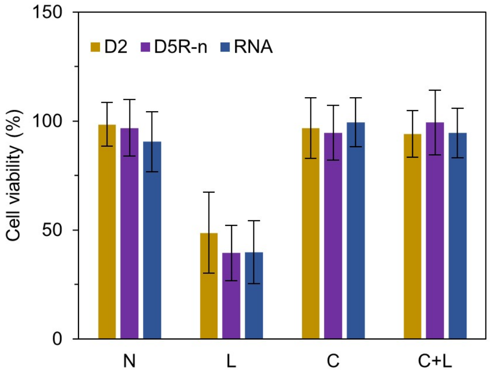


**Figure S22.** Toxicity of D2 (yellow), D5R-n (purple) and mRNA (blue) towards HeLa cells when transfected w/o transfection agent (N), using LP2000 (L), virus capsid proteins (C) or a combination (C+L).

### Note S11: Disassembly of complexed structures

Heparin can be used as a competitive agent to disassemble DNA origami complexed with virus capsid proteins, resulting in plain origami (5). A heparin molecule with an average molecular weight of 17–19 kDa consisting of on average 2.33 sulfate groups per repeating IdoA(25)-GlcNS(6S) disaccarid unit is estimated to carry a charge of -71 originating from negatively charged sulfate groups (6). The total negative charges for mRNA, D2 and D5R-n were estimated to be 996, 1626 and 1796, respectively. The amount of heparin needed is expressed as $n_{sulfates}/n_{phosphates}$. The release of D2 is monitored by the shift in electrophoretic mobility of the structure, and can be already observed at equimolar amounts (Figure S23a). To ensure the complete release, 5× excess were used for subsequent experiments.

The accessibility of the 5'-cap region after the release of mRNA-DNA origami variants from their protective virus coating was assessed. To avoid inhibiting the translation by the high salt concentration present in the complexation buffer (45 mM Tris, 75.5 mM NaCl, 10 mM acetic acid, 2.5 mM $MgCl_2$, 0.5 mM DTT and 0.5 mM EDTA) (Figure S23b, cob), the potassium acetate concentration in the translation mix was adjusted from 80 mM to 52.5 mM. It resulted in recovering the GFP signal of plain mRNA (ls, 0.0125 µg). Subsequently, complexed structures (mRNA-250, D2-250 and D5R-n-250) were treated with 5× excess of heparin for 10 min before addition to the reticulocyte lysate. A fluorescence signal could only be observed for plain mRNA (comparison to standard translation conditions, 0, *i.e.*, mRNA in RNase-free water, standard salt in translation mix).

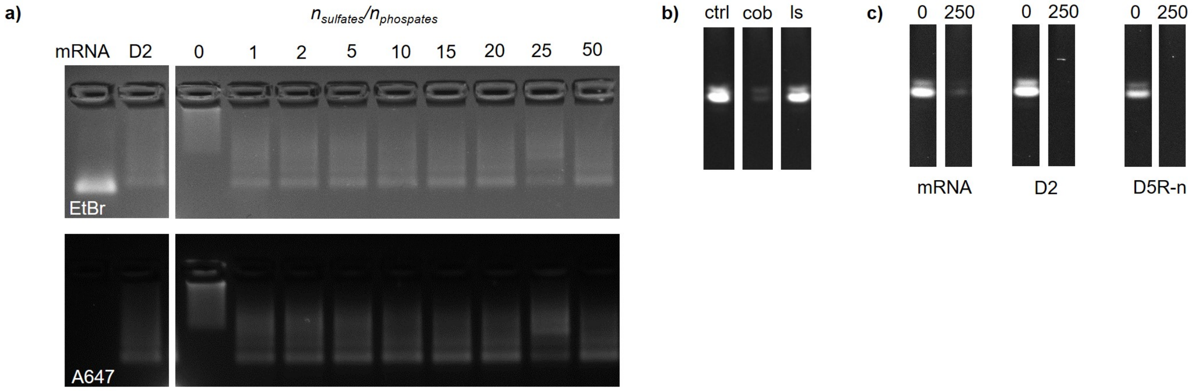


**Figure S23. a,** D2-250 can be released from its virus capsid coating by the addition of heparin. **b,** Recovery of the GFP signal from mRNA in complexation buffer (cob) by decreasing the salt concentration (ls) in the reticulocyte lysate. **c,** Translation of mRNA-250 (left), D2-250 (middle) and D5R-n-250 (right) after decomplexation with heparin using low salt conditions and comparison to the intensity of the GFP signal obtained from untreated samples (0, in RNase-free water).

### Note S12: Staple sequences for D1–D5

The staple strands to fold structures D1–D4 are listed in Tables S1–4. The design of D5 has been published previously (5), here, only the staple strands for the rerouted structure is presented (Table S5). D2 and D5R-n also list the strands facilitating the attachment of a fluorophore staple strand (A, /5ATTO590N/CAGGAACCCAAAACCG).

**Table S1.** Staple sequences for D1.

| Number | Sequence (5' to 3') |
|---|---|
| 1 | GGTAGTGCATGTGATTACGTC |
| 2 | GTAGTTGGCTCCTGGGTCAGG |
| 3 | TTGCCGTATGTTGCTTCTGCT |
| 4 | GGCGGATGCCGAGATGAACAG |
| 5 | GCCGTCCCGAGGGTGAAGAAG |
| 6 | CTCCTCGGGTAGCGGGACTTG |
| 7 | GGCACGGTGGCCGTTCGCGCT |
| 8 | GATATAGGCATGGCGCTGAAG |
| 9 | GTGGTCAAGCTCGAAACTCCA |
| 10 | CTCGCCCTGAACTTGAACTTC |
| 11 | GTGCAGATCGCCGGTTGCTCA |
| 12 | ACCTCGGTCGATGCGGGCCGT |
| 13 | ATGGTGCTACTCCAAGCTGCA |
| 14 | GCAGGACGTCGGCGGCTTGTG |
| 15 | CCCCAGGCGTCCTTGGGCCAG |
| 16 | TGGTCGGCCCTTGCTACAGCT |
| 17 | CCTTCGGACGTTGTATGTTGT |
| 18 | AAGAAGTTGCTTGTTTCACCT |
| 19 | GGGCGGACAGCACGCCTTCAG |
| 20 | CTCGATGCGCCCTCCAGGGTC |
| 21 | CGCTGCCGGTCACGCCAGGAT |
| 22 | GAACTTGGCAGCTTCTTGTAG |
| 23 | CGGCGGCGTCCTCGGGCTGTT |
| 24 | CTTGAAGCGCGGGTGCCGGTG |
| 25 | CGTCCATCTTGAAGCGGCCAT |
| 26 | CGGGCAGCTGGGTGCATCGCC |
| 27 | CGCCGATGGGGTCTACACGCT |
| 28 | TCTCGTTGGGGGTGCGTCCTC |
| 29 | GGGCACCCGCCGTAGACGTAG |
| 30 | CACTGCAACCCCGGGTGATCC |

**Table S2.** Staple sequences for D2. Strands #20–23 can be exchanged to facilitate the attachment of a fluorophore.

| Number | Sequence (5' to 3') |
|---|---|
| 1 | CGTCCATCTTGAAGCGGCCATGATATAGGCATGGC |
| 2 | GAACTTGTGGCCGTTTACGTCGCCGTCCCGAGGGT |
| 3 | GAACTTCACCTCGGCGCGGGTGCCGGTG |
| 4 | TGGTCGGCCCTTGCTACAGCT |
| 5 | ACCCCGGTGAACAGCTCCTCGGGTAGCG |
| 6 | GTGCAGATCGCCGGTTGCTCA |
| 7 | CGGCGGCGGTCACGCCAGGATGGGCACC |
| 8 | CGGGCAGCTGGGTGCATCGCC |
| 9 | GGGCCAGGGCACGGGCAGCTTCTTGTAG |
| 10 | CTCGCCCTGAACTTCAGGGTC |
| 11 | GTGGTCAAGCTCGAAACTCCA |
| 12 | GGGCGGACAGCACGCCTTCAGCTCGATGCGCCCTC1 |
| 13 | GTCCTCGGGCTGTTGTAGTTG |
| 14 | GGACTTGAAGAAGTTGCTTGTTTCACCT |
| 15 | GCAGGACGTCGGCGGCTTGTGCCCCAGGCGTCCTT |
| 16 | TTGCCGTATGTTGCTTCTGCTGGTAGTGCATGTGA |
| 17 | GAAGAAGATGGTGCGCTCCTGGGTCAGG |
| 18 | TTGAAGTCGATGCGGGCCGTC |
| 19 | TACTCCAAGCTGCACGCTGCC |
| 20 | TCGCGCTTCTCGTTGGGGTCTACACGCT |
| 21 | GCCGATGGGGGTGCGTCCTCC |
| 22 | GCTGAAGCACTGCACGCCGTAGACGTAG |
| 23 | CCTTCGGACGTTGTATGTTGTGGCGGATGCCGAGAGTGATCC |
| 20-A | TCGCGCTTCTCGTTGGGGTCTACACGCTTTCGG TTT TGG GTT CCT G |
| 21-A | GCCGATGGGGGTGCGTCCTCCTTCGG TTT TGG GTT CCT G |
| 22-A | GCTGAAGCACTGCACGCCGTAGACGTAGTTCGG TTT TGG GTT CCT G |
| 23-A1 | CCTTCGGACGTTGTATGTTGT |
| 23-A2 | GGCGGATGCCGAGAGTGATCCTTCGG TTT TGG GTT CCT G |

**Table S3.** Staple sequences for D3.

| Number | Sequence (5' to 3') |
|---|---|
| 1 | GCCGCTTTTCTCGTTGGGGTCTTTGCTC |
| 2 | TGTTGTGGCGGATCCGTCGCC |
| 3 | TGAACAGCTCCTCGGGTGGTCTGTAGTT |
| 4 | CGAGAGTACACGCTGAACTTGTGGCCGT |
| 5 | CAAAGACGTGTTCTGCTGGTAGTGGTCGGC |
| 6 | ATCGCGCACTTGTACAGCTCG |
| 7 | AGGGCGGATATAGACGTTGTGCCCTCGAACTTCAC |
| 8 | GGCAGCTTGCCGGTAGCTCGATTAAGCG |
| 9 | GGTGCGCTCCTGGAGGGGTAGCGGCTGAAGACTGC |
| 10 | GCTCAGGAAGGCAAGCCCCGCAGAAGGC |
| 11 | AAGAAGTCGTGCTGCTTCACTTTTCTCTTTATT |
| 12 | TGTTGCCAAGTCGATGCCCTTCAGCTCG |
| 13 | GGTGTCGGCTGTTGTAGTTGT |
| 14 | GCCGTCGGATGCCGTTCTTCT |
| 15 | GATGGGGCAAGAGGTACAGGTGCAAGGG |
| 16 | AGAGAAGTCACCATGGTGGCTCTTATAT |
| 17 | TTCTTCTTACTCTTTGTGGTCCGTAGCC |
| 18 | CTTCCTACTCAGGCTCTTATTTCCTC |
| 19 | ACGGGGCTTGAAGTTCACCTTTCCTTGAAGAAGAT |
| 20 | CCTTGGTCCT |
| 21 | CGCCCTCGCCGGGATCCCGGC |
| 22 | GCTTGTCGGCCATGACTGGGT |
| 23 | ACTCCAGCTTGTGCCCATGTG |
| 24 | GGCGGTCACGAACTCC |
| 25 | ATGCGGTATGAACTTCAGGGTCAGCTTGCCGTAGGTGGCATCGCCCT |
| 26 | TTACGTCGCCGTCCGGTGCAGTCACCAG |
| 27 | AGCTTAACCAGGATGGGCACCACCCCGG |
| 28 | ACGCCGTAGGTCAGCCCTTGCAAGGGCA |
| 29 | CTCGGCGCGGGTCTACGAGGGTGGGCCAGGGCACG |
| 30 | GAGCTGCACATGGCGGACTTG |
| 31 | TTCGGGCGCTGCCGTCCTCGA |
| 32 | TGGCCAGTAGTGGTTGTCGGGCAGCAGC |
| 33 | TCCATGCAGCAGGACCCAGGA |

**Table S4.** Staple sequences for D4.

| Number | Sequence (5' to 3') |
|---|---|
| 1 | AGGACCATGTGATCGCTTACTTGTACAGCTCGTCC |
| 2 | TCGATGCCCTTCAGCCAGGGCTGGCATC |
| 3 | GTCACGAGGGTGGGCTCGATGTTAATTA |
| 4 | CGATGGGGGTGTTCTGCTGGTTGGTTGTCGGGCAGCAGCACG |
| 5 | AGGGAGAGTCTTGTAGTTGCCGTCGTCC |
| 6 | TCGAACTTCACCTCTGAAGACCGTTTAC |
| 7 | GCCCTCGCCCTCGCCGGACACGCTGAACTGCACGC |
| 8 | CAGATAGTTGTACTCCCCAGGGAGTGATCC |
| 9 | GGCGAGCTTGTGGCTGCACGCCGTAGGTCAGGGTG |
| 10 | CTTATATACCACCCGAAGTCGTGCTGCTTCATGTG |
| 11 | ATGCCGAATGTTGCCGTCCTCCTTGAAG |
| 12 | TGCCGTCCTCGATGTTGTGGCTCGTTGGGGTCTTTGCTCAGG |
| 13 | CTTCCTACTTGGACGTAG |
| 14 | GGGCCCTTCTTACTAGACCAAGAGGTACAGGTGCA |
| 15 | CCTTCGGGCATGGCGGGCGCTCCCAGGCT |
| 16 | TTGAAGAAGATGGTACTTGAACGGTGAACAGCTCCTCGCC |
| 17 | GGCATGGCAGGTAGAGTGGTC |
| 18 | AGGCAGCCGGTTCACCAGGGTGTCGCCC |
| 19 | AGACGTTGTGGCTGTT |
| 20 | CCATGATATC |
| 21 | TGCCGTTCTTCTGCTTGTCGGGCGGCGGTCACGAACTCCAGC |
| 22 | GTCGCCGTCCAGCTCGACCAGGATGGGCTTGTCGC |
| 23 | GCGGACTGGGTGCTCCAGAAGGCAAGCCCCGCAGA |
| 24 | GAAGTTCCCGTAGGACGGGCAGCTTGCCGGTGGTG |
| 25 | AGCGGCCGCGCTTCGGATCTT |
| 26 | TTGTGCCAGC |
| 27 | GTCGGGGTAGCGGCGGCGCGGGAAGAAG |
| 28 | GTGAACTTCAGGGTCAGCTTGACCTTGA |
| 29 | CTTGCTCACCATGGTGGCT |
| 30 | TTATTCAACTTCTTTTCTCTCTTATTTCCTC |

**Table S5.** Staple sequences for D5R-n. Strands 1–18 represent the original strands for the core of the structure, #19–22 can be exchanged to facilitate the attachment of a fluorophore, and #28–29 are the two replacement stands folding the 5'-cap region. They can be exchanged to contain a 6 nt or a 10 nt toeholds, so the replacement strands can be removed from folded structures by the addition of the respective invader strand pair.

| Number | Sequence (5' to 3') |
|---|---|
| 1 | TGGTCGGACGCTGAGGTGGGCCAGGGCACCGCCCTCG |
| 2 | TTGCCGGTGGTGCAGATGAACT |
| 3 | TTGATGCCGTTCTTCTGCTTG |
| 4 | ACTTGAAGAAGTCGTGCTGCTTCAT |
| 5 | TTTGCTCAGGGCGGAGCAAGCCCCGCAGAAGGCAGCT |
| 6 | GAAGATGGTGCGCTCCTGGACGCTTTATTC |
| 7 | CCGGCGGCGGTCACGAACTCCA |
| 8 | TAGGTGGCATCGCCCTCGCCCTCGCCGGACCGAGCTGC |
| 9 | GGATGTTGCCGTCCTCTCCAGCTTGTGCCCCA |
| 10 | GTGGTCGGGACCACCCCGGTGAACAGCTC |
| 11 | TCGGCCATGATATAGACGTTGTGGCTGTTCACC |
| 12 | TCAGGGTAGTTGTACCTTGAAGT |
| 13 | ACGCTGCCGTCCTCGATGTTGTGATCGCGCTTCTCGTTGGGGTC |
| 14 | CTTCCTACTCAGGTAGCCTTCGGGCATGGCGG |
| 15 | CCAGAAGCTGGGTGCTGGTAG |
| 16 | TTACTTGACCATGTGGCGGAT |
| 17 | GCAGGTACAGCTCGTCCATGCCGAGAGTGATC |
| 18 | TAATTAAGTTCACCAGGGTGTGGGCAGC |
| 19 | AACTTCACCTCGGCGCGGGTCTGGGCATGG |
| 20 | CGTAGGTCAGGGTGGTCACGAGACTTGTG |
| 21 | CTTGAAGTTGTCAGCTTGCCG |
| 22 | CGATGCCCTTCAGCTCGATGCGGCGGCCGC |
| 23 | GCCGTTTACGTCGCCGTCCAGCTCGACCAG |
| 24 | GTGGTTGAGAAGAATGTAGTTGCCGTCGTGCACGC |
| 25 | GCCGTCGCCGATGGGGGTGTTCTGCTCAGGTA |
| 26 | AAAGACCAAGAGGTACAGGTGCAAGGGAGTCGGGCA |
| 27 | GATGGGCGTAGCGGCTGAAGACTCCTTGAA |
| 28 | CTCGCCCTTGCTCTCTTTTCTCTCTTATTTCCTC |
| 29 | GCAGCATATTTCTTCTTACTCTACCATGGTGGCTCTTACGGG |
| 19-A | AACTTCACCTCGGCGCGGGTCTGGGCATGGTTCGG TTT TGG GTT CCT G |
| 20-A | CGTAGGTCAGGGTGGTCACGAGACTTGTGTTCGG TTT TGG GTT CCT G |
| 21-A | CGATGCCCTTCAGCTCGATGCGGCGGCCGCTTCGG TTT TGG GTT CCT G |
| 22-A | CTTGAAGTTGTCAGCTTGCCGTTCGG TTT TGG GTT CCT G |
| 28-6 | CTCTCCCTCGCCCTTGCTCTCTTTTCTCTCTTATTTCCTC |
| 29-6 | CGGACGGCAGCATATTTCTTCTTACTCTACCATGGTGGCTCTTACGGG |
| 28-10 | TCTCTCCCTTCTCGCCCTTGCTCTCTTTTCTCTCTTATTTCCTC |
| 29-10 | ACGGACGCAAGCAGCATATTTCTTCTTACTCTACCATGGTGGCTCTTACGGG |
| inv-6(28) | GAGGAAATAAGAGAGAAAAGAGAGCAAGGGCGAGGGAGAG |
| inv-6(29) | GAGGAAATAAGAGAGAAAAGAGAGCAAGGGCGAGAAGGGAGAGA |
| inv-10(28) | CCCGTAAGAGCCACCATGGTAGAGTAAGAAGAAATATGCTGCCGTCCG |
| inv-10(29) | CCCGTAAGAGCCACCATGGTAGAGTAAGAAGAAATATGCTGCTTGCGTCCGT |

### Note S13: Estimation of DNA origami concentration

The concentration of mRNA-DNA origami variants was estimated using Lambert-Beer's law. The variant-specific extinction coeffienct was estimated from the number of hybridized and non-hybridized nucleotides (7).

**Table S6.** Extinction coefficient of different mRNA-DNA origami variants

| mRNA-DNA origami variant | Extinction coefficient ($M^{-1}$ $cm^{-1}$) |
|---|---|
| D2 | $1.21 \times 10^7$ |
| D5 | $1.29 \times 10^7$ |
| D5R-n | $1.27 \times 10^7$ |
| D5R-6 | $1.30 \times 10^7$ |
| D5R-10 | $1.304 \times 10^7$ |
